# Exploring the role of stimulus similarity on the summation effect in causal learning

**DOI:** 10.1101/178954

**Authors:** Omar D. Pérez, Rene San Martín, Fabián A. Soto

**Author notes:** Address correspondence to: Omar D. Pérez, Division of Humanities and Social Sciences, California Institute of Technology, Pasadena, California, CA 91125.

## Abstract

Several contemporary models of associative learning anticipate that the higher responding to a compound of two cues separately trained with a common outcome than to each of the cues alone -a summation effect-is modulated by the similarity between the cues forming the compound. Here, we explored this hypothesis in a series of causal learning experiments with humans. Participants were presented with two visual cues that separately predicted a common outcome and later asked for the outcome predicted by the compound of the two cues. Importantly, the cues’ similarity was varied between groups through changes in shape, spatial position, color, configuration and rotation. In variance with the predictions of these models, we observed similar and strong levels of summation in both groups across all manipulations of similarity (Experiments 1-5). The summation effect was significantly reduced by manipulations intended to impact assumptions about the causal independence of the cues forming the compound, but this reduction was independent of stimulus similarity (Experiment 6). These results are problematic for similarity-based models and can be more readily explained by rational approaches to causal learning.

---

In natural environments, widely different stimuli are grouped together in classes (e.g., plants, cats, birds, etc.) and related with similar outcomes (e.g., nutrition, danger of injury or death, a potential mate, etc.). Encoding these relationships is critical for organisms, as it allows them to take advantage of the regularities in the environment. Consider, for example, a foraging animal that has got mildly sick after eating a particular food. The more the animal is able to estimate the level of sickness produced by foods with similar smell or coloration, the higher will be its chances of surviving and reproducing. This process, known as *generalization*, allows the animal to extract valuable information from past events and apply it to novel situations (Brandon, Vogel, & Wagner, 2000; Rescorla, 1976; Soto, Gershman, & Niv, 2014b; Soto & Wasserman, 2010). By generalizing between current and past events, organisms learn to respond similarly to different stimuli.

A particular form of stimulus generalization, known as *compound generalization*, is also critical for organisms to thrive in natural environments. The most typical example of this type of generalization is the *summation effect*, where the animal needs to estimate the level of sickness produced in the eventual case of eating two types of food that have independently produced a mild level of sickness in the past. From a psychological standpoint, the question at issue is whether the animal will predict a similar level of sickness for the compound of the two foods than to each of the two foods alone, or a stronger level of sickness for the compound of the two foods than to each of the two foods alone, after having learned the causal status of each one the foods in causing the sickness. If the prediction is that the level of sickness will be stronger, then it is said that summation has occurred.

In the lab, the summation effect is studied through a design in which two different stimuli, A and B, are separately paired with a common outcome and subsequently tested in compound (AB) (Collins & Shanks, 2006; Soto, Vogel, Castillo, & Wagner, 2009). A summation effect is obtained in this context if the response to the compound AB is greater than the response to the components A or B presented separately.

Different models of Pavlovian conditioning make different predictions for summation studies. These models have followed two theoretical traditions, each with a different view on how the stimuli A and B are represented separately and in compound. On the one hand, elemental models (e.g., Rescorla & Wagner, 1972) claim that a stimulus is represented by a finite (potentially large) number of elements which separately acquire their own associative strength in predicting the outcome. In this theory, the summation effect is anticipated under the assumption that the associative strengths accrued by A and B should sum linearly when presented in compound.

A fundamentally different view, known as *configural theory*, was proposed by Pearce (1987, 1994), who claimed that subjects would represent the compound AB as a different stimulus (say AB=C) during testing. According to Pearce, responding to the compound AB (i.e., C) will depend upon the degree to which the original learning to A and B generalizes to this new stimulus representation - that is, the extent to which the compound AB is considered to be similar to the individual stimuli A and B. From a pure configural perspective, half of the associative strength from each component is generalized to AB: responding to AB should thus be similar to the average of responding to A or B alone.

The evidence regarding the summation effect in Pavlovian conditioning is mixed. Although some studies have found the effect (Myers, Vogel, Shin, & Wagner, 2001; Rescorla, 1997; Whitlow & Wagner, 1972) others have failed to report it (Aydin & Pearce, 1994, 1995, 1997; Rescorla & Coldwell, 1995). Several authors (see, for example, Wagner 2003; Soto et al. 2014) have interpreted these results as arising from the different modalities of the stimuli used in each study. Indeed, failure to report summation is usually associated with the employment of stimuli from the same modality, whereas reliable summation is usually observed in studies that have employed stimuli from different modalities. Using rabbit eyeblink conditioning, for example, Kehoe and colleagues (1994) reported higher responding for a group trained with tone and light components than in a group trained with tone and noise components. Similar results were obtained more recently by Thein and colleagues (2008) using rats’ magazine approach.

Several contemporary theories have interpreted these results as being a consequence of the perceptual interaction of same- and different-modality components (Harris, 2006; Soto, Gershman, & Niv, 2014; Thorwart, Livesey, & Harris, 2012; Wagner, 2003). In particular, these theories claim that high similarity of components, typical of unimodal stimuli, should encourage more configural processing; in contrast, the lower similarity of multimodal components should foster elemental processing. A weak summation effect should therefore be observed when the components A and B are similar, whereas stronger levels of summation should come about when these components are dissimilar (Harris & Livesey, 2010; Harris, 2006; Kinder & Lachnit, 2003; Soto et al., 2014; Soto, Quintana, Pérez-Acosta, Ponce, & Vogel, 2015; Thorwart et al., 2012; Wagner, 2003). The mechanisms underlying this prediction vary among models, but the idea of similarity as a critical factor is widely present. Using an elemental approach, for example, Wagner’s (2003, see also Brandon et al., 2000) Replaced Elements Model (REM) allows for some processing flexibility, by allowing the representation of two separate stimuli A and B to be partially “replaced” by a unique representation when they are presented in the compound AB. Critically, the model assumes that the level of replacement is a function of the similarity of A and B. This notion allows the model to be flexible enough with respect to generalization strategies, anticipating higher summation for dissimilar components (Brandon, 2003; Wagner, 2003, 2008).

Studies in human causal learning have also demonstrated that stimulus properties mediate the type of representation employed by subjects and impact on several learning phenomena. In a blocking scenario, for example, participants experience A followed by an outcome over a number of trials and are subsequently presented with the compound AB followed by the same outcome over a comparable number of trials. The fact that the causal rating for B is usually lower than the rating for A is regarded as evidence of cue A *blocking* the acquisition of causal strength to B (Dickinson, Shanks, & Evenden, 1984). In an experiment reported by Livesey and Boakes (2004, Experiment 3), a compound of two stimuli combined into a single object significantly reduced blocking to B compared to a condition in which the two cues were presented separated on the screen. Presumably, the stimuli presented as a single object were processed as a single configuration, which reduced any competition between individual cues to be associated with the outcome. The opposite effect occurs when elemental processing is encouraged by presenting the cues in different positions on the screen: under this condition, blocking is more likely to be observed (Glautier, 2002; Livesey & Boakes, 2004). Given these results, contemporary theories have incorporated stimulus properties such as spatial position and organization as another aspect of stimulus similarity (Soto et al., 2014a; Thorwart et al., 2012). For example, Soto et al.’s (2014) model combines the notion of consequential regions proposed by Shepard (1987) with contemporary Bayesian approaches to generalization (Courville, 2005; Courville, Daw, & Touretzky, 2006) and makes the same prediction as flexible associative models: similar stimuli should produce a weaker summation effect than dissimilar stimuli. Importantly, Soto et al. argue that factors such as spatial contiguity should be critical in determining whether summation should be obtained. A similar role for spatial contiguity can be found in Harris and Livesey’s (2010; see also Thorwart et al., 2012) model, who propose that the closer two stimuli are, the more their representations interact with each other (through normalizing gain control), producing configural processing. In the rest of this paper, we use the term similarity specifically in regard to all the manipulations of stimulus properties that are considered by these models to mediate the generalization process: more similarity will thus indicate more configural processing; less similarity more elemental processing.^1^

Summation in human causal learning shows some parallels to the results reported in Pavlovian conditioning procedures. For example, multimodal stimuli seem to produce higher summation than unimodal stimuli (Redhead, 2007). However, the summation effect has not been studied, neither in human causal learning nor in animal conditioning, employing unimodal components with varying degrees of similarity. This is the goal of the series of studies reported in this paper.

To test whether manipulating similarity of visual cues can exert an effect on summation, we adapted the widely-used “allergy prediction” task (Van Hamme & Wasserman, 1994) where participants are trained to predict allergies in a hypothetical patient. In our version of this task, participants were asked to predict allergies to drugs which were represented by visual shapes. During testing, participants had to predict the outcome produced by compounds of these cues. The critical manipulation involved changing properties of the cues to make them more similar or dissimilar. In particular, the shapes of the cues varied in configuration, rotation, spatial position, and color, factors which allowed us to easily manipulate the properties of stimuli that contemporary models consider to be critical in promoting configural or elemental processing. The design included two groups of subjects. In one of the groups (group *intra*, i.e., intra-dimensional difference), cues A and B differed by some degree within a single dimension while in the other they differed across two or more dimensions (i.e., extra-dimensional difference; group *extra*)^2^

Thus, unlike previous experiments exploring the extent to which stimulus properties would impact on the *learning process* and hence modulate responding to the components because of the accruement of different associative strengths (see, for example, Livesey & Boakes, 2004), our experiment varied the degree of similarity of visual cues to test whether this factor would have an effect on the *generalization strategy* after the learning stage, bringing about different levels of summation in groups *extra* and *intra*. Several contemporary learning theories predict a stronger summation effect in group *extra* than in group *intra*, both when these stimuli vary in featural similarity (Harris, 2006; Harris & Livesey, 2010; McLaren & Mackintosh, 2002; Soto et al., 2014; Soto et al, 2015; Thorwart et al., 2012; Wagner, 2003) and when they vary in others stimulus factors such as spatial separation (Harris & Livesey, 2010; Soto et al., 2014; Thorwart et al., 2012). The resulting data should allow as to probe the theoretical assumption underlying these models that stimulus similarity in general, and not stimulus modality in particular, is what controls compound generalization.

## Experiment 1

The stimuli used for cues A and B in Experiment 1 are shown in Figure 1. Cues in group *extra* varied in both shape and color, whereas cues in group *intra* varied only in shape.

**Figure 1.**
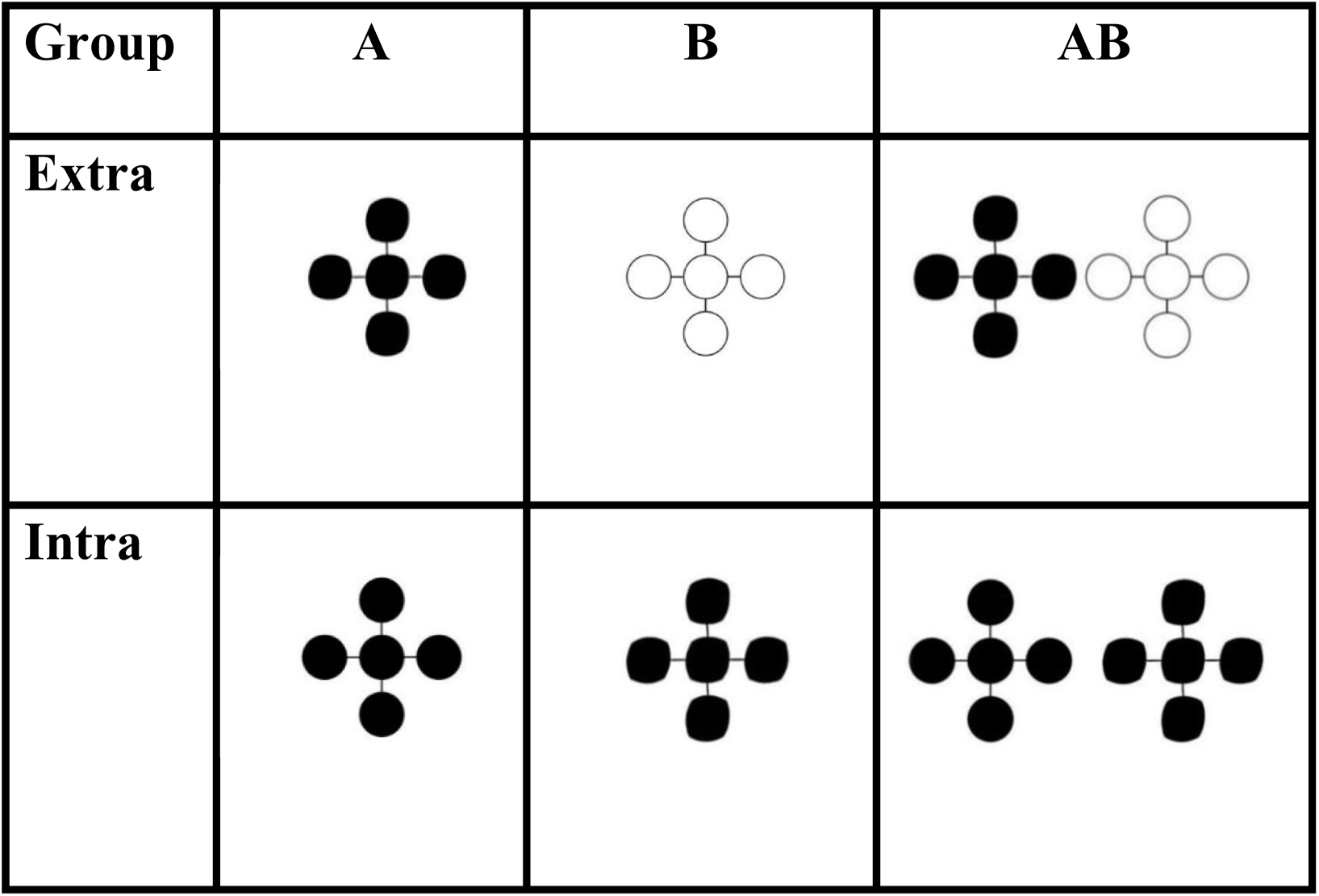
Target stimuli used in Experiment 1. Minimal variations in shape were used in both groups. Maximal variations in color were used in addition for group *extra*. The positions of stimuli A and B in the compound AB were counterbalanced during both training and testing.

Table 1 shows the design of this experiment. Training consisted of two different cues which independently predicted allergic reaction (A+/B+) and two fillers which independently predicted no allergic reaction (C-/D-). There were two further compound fillers which predicted allergic reaction (EF+) and no allergic reaction (GH-). On training, participants assessed whether the allergy would be present or not by using a binary response (allergy/no allergy). Feedback was provided about the magnitude of the allergic reaction produced by the cue. During the test stage, participants had to estimate the magnitude of allergic reaction for different cues and combinations of cues, by entering it on a rating scale.

**Table 1.** Design of Experiment 1. The only difference between the group intra and *extra* was the dimensions in which the stimuli A and B differed during training.

| Training | Test |
| --- | --- |
| **Stimuli** | A |
| A+ | B |
| B+ | EF |
| EF+ | AB |
| **Fillers** | C |
| C- | D |
| D- | GH |
| GH- | CD |

*Note:* Letters represent different chemical shapes which could be followed (+) or not followed (-) by allergic reaction in a hypothetical patient.

Similarity-based models predict that the relatively similar stimuli shown to group *intra* should be processed more configurally and therefore promote lower summation than the relatively dissimilar stimuli shown to group *extra*, which should be processed more elementally and therefore promote higher summation.

## Method

### Participants

30 undergraduate students from the University of Santiago, Chile. Participants were randomly assigned to one of two groups (*n*=15 each). All participants were tested in the same room at the same time and did not have any previous experience with similar research. They were compensated with course credit for their participation. All procedures were approved by University of California Santa Barbara IRB Protocol 13-0425: *The Cognitive Neuroscience of Human Category Learning*.

### Materials

Participants were tested on Intel i7 computers running Psychopy 1.77 (Peirce, 2007) under Windows 7. The experiment was programmed with Psychopy 1.75. Computers were connected to 17-inch Samsung color monitors.

### Procedure

At the beginning of the experiment, participants were informed that all the instructions would be presented on the screen. They were randomly assigned to different computers and started the experiment at the same time.

The only difference between groups *intra* and *extra* was the stimuli assigned to the roles of A and B in the design. For group *intra*, A and B varied along a single stimulus dimension (shape), whereas for group *extra*, A and B varied along two stimulus dimensions (color and shape, see Figure 1). During each trial of the training phase, a stimulus or pair of stimuli were presented at the top-center of the screen, followed by the phrase “press ‘a’ to indicate an allergic reaction, and ‘n’ to indicate no allergic reaction” at the bottom of the screen. The stimuli were presented in one of two positions: immediately to the left of the horizontal middle line of the screen, or immediately to the right of that line. Because all the theories we are testing anticipate that the assignment of the stimulus to left or right position in each trial will be irrelevant for the generalization process, the position of each cue was randomly chosen during each trial.

After the participant entered a response, feedback was provided at the bottom-center of the screen for 2 seconds. On positive trials (marked + in design table) feedback consisted of the phrase “Allergic reaction” in red, followed in the next line by the phrase “10 points of allergy over a total of 20.” On negative trials, feedback consisted of the phrase “No allergic reaction” in black, followed in the next line by the phrase “0 points of allergy over a total of 20.”

Participants completed 25 blocks of training, each consisting of a single presentation of each of the 6 trial types shown in Table 1. Trial types were presented in random order within each block; the assignment of stimuli to the A-H conditions was semi-random. There were two pairs of stimuli that could be assigned to the roles of A and B. The stimuli in each pair were chosen based on similarity ratings obtained in a pilot study, so that the dissimilarity between A and B in group *extra* was higher than the dissimilarity between A and B in group *intra*. There were 6 other stimuli that differed in both shape and color and that were randomly assigned to the roles of C-H for each subject. During the testing phase, a new set of instructions was presented to the participants and then each of the trial types shown in the right column of Table 1 was presented once. The test stimulus was presented at the top-center of the screen, followed by the instructions “estimate the level of allergic reaction that this drug will cause in Mr. X.” At the bottom of the screen, there were 21 circles with numbers inside ranging from 0 to 20. The left side of the scale was labeled “no allergic reaction” and the right side of the scale was labeled “maximal allergic reaction.” No feedback was presented during the test stage.

## Results and Discussion

For all the experiments reported in this paper, the following criterion was used to exclude participants from the main analysis: All participants that failed to score on average 10 (+/- 3) points of allergy to cues that predicted allergy and 0 (+3) points of allergy to cues that did not predict allergy, were left out from the analysis. The rationale behind this criterion was to include data only from participants who successfully learned to discriminate between cues that predicted allergy and those that did not predict allergy.

The statistical analyses for all the experiments reported in this paper were performed using R version 3.3.0 and RStudio version 1.0.136 (RStudio Team, 2015), as follows. After fitting a linear-mixed model using the library nlme version 3.1-131 (Pinheiro, Bates, DebRoy, Sarkar, & R Core Team, 2016), a 2 x 3 mixed analysis of variance (ANOVA) was run. The ANOVA distinguished one between-subjects factor (group, with two levels: intra and extra) and one within-subjects factor (cue, with three levels: A, B and AB). A rejection criterion of *α* = .05 was used. When the *F* test yielded a detectable difference for any factor or interaction, a Tukey’s HSD (honest significant difference) post-hoc test was performed on those factors, using Bonferroni’s method to correct for multiple comparisons. Eta- squared (*η*^2^), along with a 90% confidence interval (CI) on this estimate, was calculated as a measure of effect size for each factor and interaction. A summation effect would be revealed in the post-hoc comparisons if the compound AB is scored higher than each of A and B. Different summation effects between the groups would be revealed by a significant Cue x Group interaction.

Images with all the stimuli used in the task and the data obtained in the experiments reported in this paper can be found online at github.com/omadav/similarity_summation.

Thirteen participants did not meet the criterion in this study. The final number of participants per group was *n*_*intra*_ = 9, *n*_*extra*_ = 8. The results are shown in Figure 1. Both groups reported higher scores for the compound than for the individual cues (F_2, 30_=907 .8, p=.00, *η*^2^=.98 90% CI [.97, .99]), but no effects of group were found (F_1, 15_=1.95, p=.18, *η*^2^=.11 90% CI [.00, .36]). Importantly, there was no detectable difference in summation between the groups, as revealed by a nonsignificant interaction between group and cue (F_2, 30_=1.74, p=.19, *η*^2^ =.10 90% CI [.00, .25]). Post hoc analysis revealed significant differences for cues A and AB, and for cues B and AB (p<.01 in both cases).

These results show that summation can be obtained in causal learning when the stimuli forming the compound differ either in color and shape (group *extra*) or only in shape (group *intra*). This expands on previous reports of summation in human causal learning (Collins & Shanks, 2006; Glautier, Redhead, Thorwart, & Lachnit, 2010; Soto et al., 2009; Vadillo, Ortega-Castro, Barberia, & Baker, 2014) and attests to the generality of this effect. However, there was no effect of stimulus similarity on the magnitude of summation obtained.

To test whether participants were rating compounds unconditionally higher than single stimuli (i.e., irrespective of the experience during training), we compared the rating obtained for AB with that for EF. The analysis showed that there was a significant difference between AB and EF in group intra (mean difference=8.83 95% CI [7.05, 10.61], t(8)=11.43, p<.001, D=3.81 95% CI [1.86, 5.74]). In group extra, all participants scored AB as producing 20 points of allergy and EF as producing 10 points of allergy; this uniformity of results prevented us from performing a statistical test of their difference.

As Figure 1 shows, both groups scored the compound as being close to the linear sum of the individual predictions (around 20 points of allergy). It is possible that this is a consequence of the employment of an allergy scale which prompts participants to score the compound as the exact sum of the two individual predictions (the maximum possible rating was exactly double the magnitude of allergy predicted by cues A and B during training). In Experiment 2, we addressed this potential shortcoming of the design by using an allergy scale with a maximum level of allergy that was higher than 20 points. In addition, a different set of stimuli was used, varying in the dimensions of color and rotation, to increase the difference between group in similarity of A and B.

## Experiment 2

The same design as in Experiment 1 was used in this experiment, except that the target stimuli now varied in color and rotation. Having established the presence of a strong and clear summation effect with our task in Experiment 1–which included comparisons of AB with A, B, and control compounds EF and CD–all following experiments focus on testing the effect of similarity on the summation effect as evaluated through the simple comparison of AB and its components A and B. The design, therefore, included A and B as targets, and C and D as fillers, but excluded other compounds and fillers from the previous design (see Table 2). The shape was now the same in both groups. In group *intra*, stimuli A and B were made near identical, by having them differ slightly in rotation, whereas in group *extra*, A and B were made sharply different, by having them differ not only in rotation but also in color (see Figure 3).

**Table 2.** Design of Experiments 2, 3, 4 and 5. The only difference between the group intra and *extra* was the dimensions in which the stimuli A and B differed during training.

| Training | Test |
| --- | --- |
| **Stimuli** | A |
| A+ | B |
| B+ | AB |
| **Fillers** | C |
| C- | D |
| D- |  |

**Figure 2.**
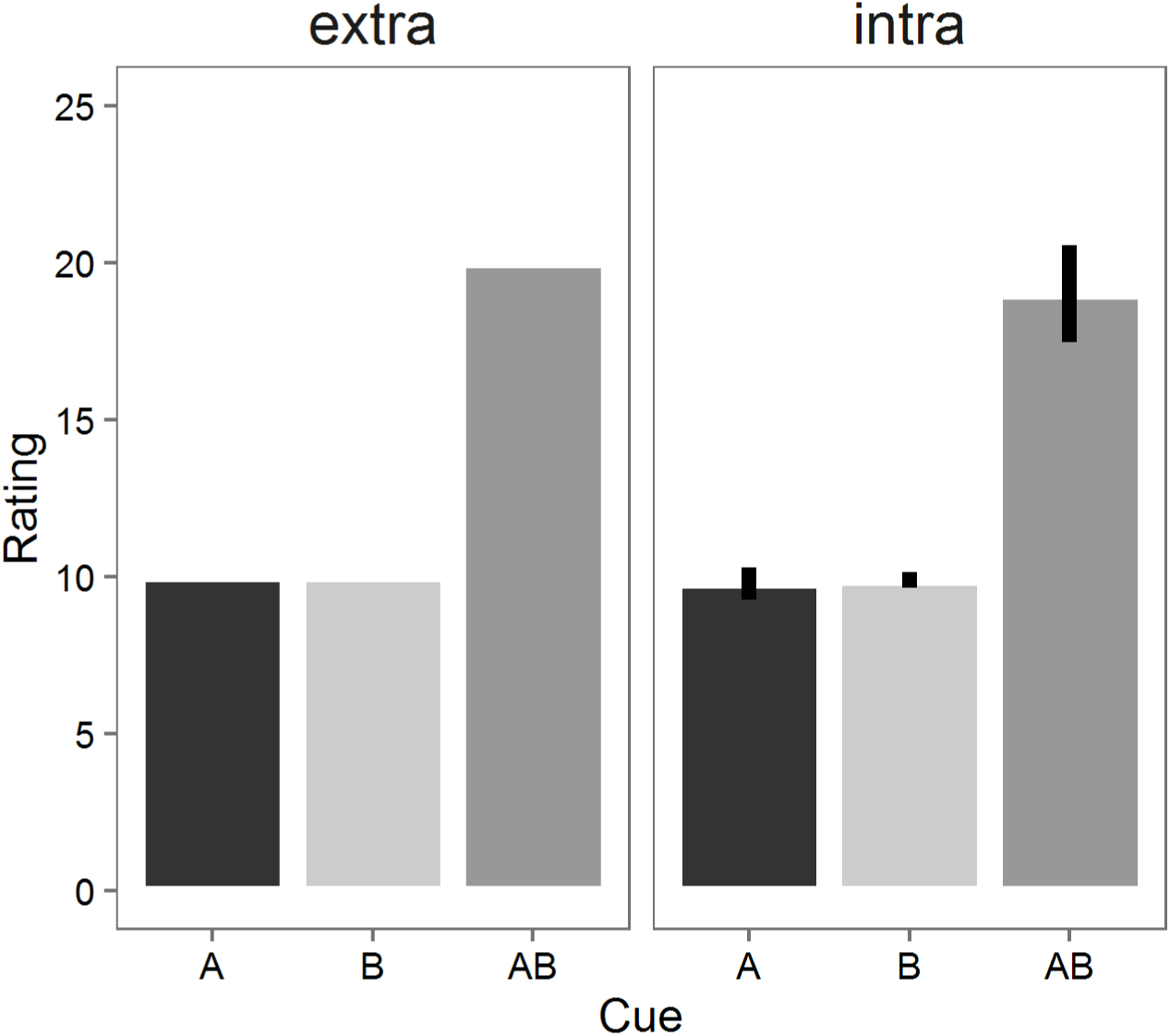
Mean causal ratings for cues A, B and AB for groups *extra* and *intra* in Experiment 1. Error bars represent 95% confidence intervals.

**Figure 3.**
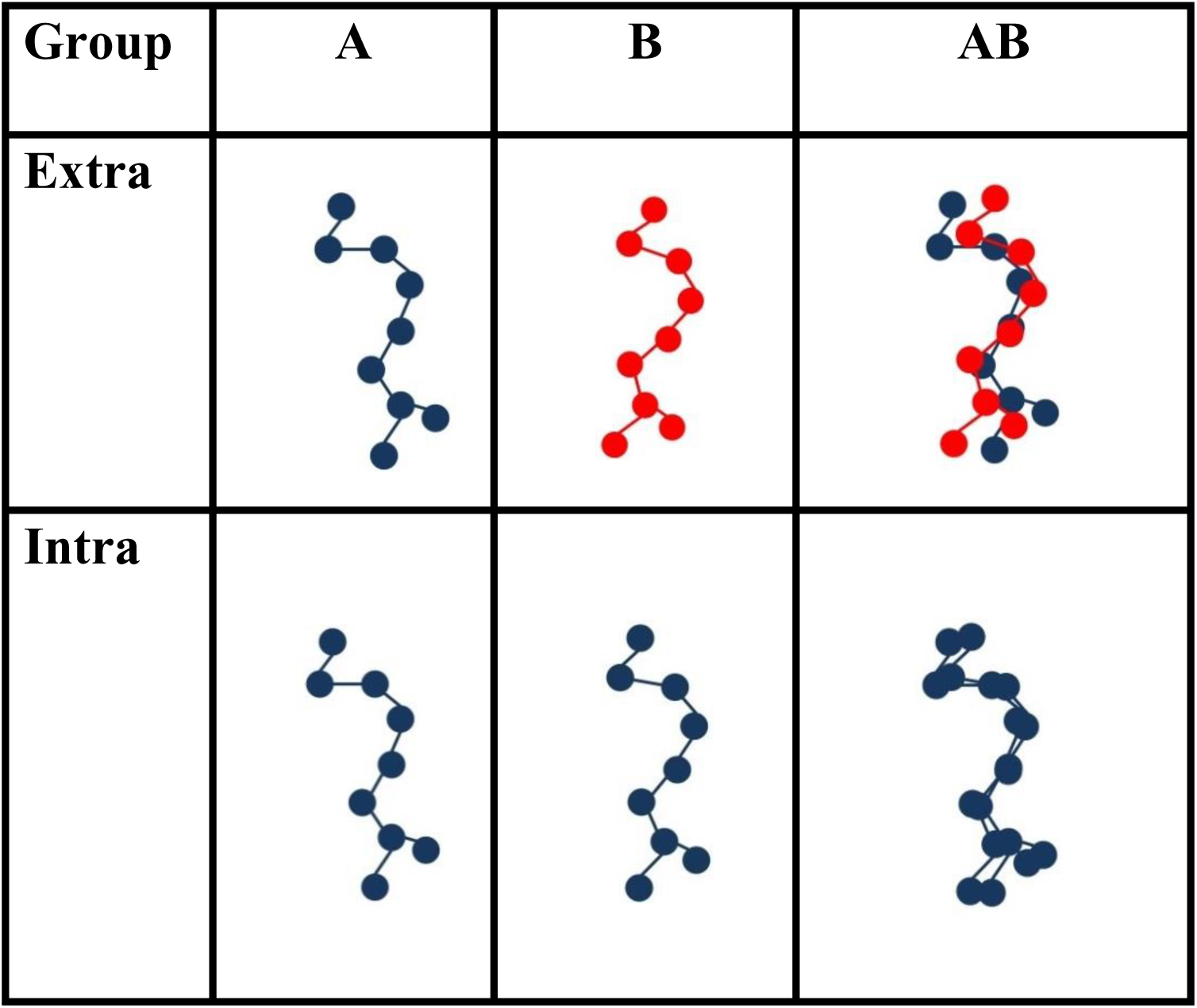
Target stimuli used in Experiment 2. Minimal variations in rotation were used in both groups. Variations in color were used in addition for group *extra*.

In Experiment 1 there was a very strong summation effect in both groups. From an associative perspective, this implies that both groups were deploying strongly elemental strategies in forming their predictions. For example, cues were presented in clear spatial separation, and previous data suggest that spatial separation is a critical factor in modulating configural and elemental processing (Livesey & Boakes, 2004; Melchers, Shanks, & Lachnit, 2008; Soto et al., 2014b; Thorwart et al., 2012); spatially contiguous stimuli, in contrast, should foster configural processing (Glautier, 2002; Livesey & Boakes, 2004; Melchers et al., 2008). To encourage configural processing in both groups in Experiment 2, both A and B were presented at the center of the screen.

In addition to changes in stimulus properties, the feedback during training was modified so that the maximum possible outcome was 35 points of allergy. The value was explicitly chosen to be higher than the sum of the outcomes of each of the individual cues A and B – we reasoned that using a maximum of 20 points of allergy could have prompted participants in Experiment 1 to score the compound as the sum of the two individual components. The rating scale during testing was also changed accordingly.

## Method

### Participants

38 undergraduate students from the University of California Santa Barbara. Participants were randomly assigned to one of two groups (*n*=19 each). They were tested in the same way as in Experiment 1. Participants were compensated with course credit for their participation.

### Materials

Participants were tested on Macintosh computers running Psychopy 1.77 (Peirce, 2007) under OSX. The experiment was programmed in Psychopy 1.75.

### Procedure

The procedure was the same as in Experiment 1. The main differences were the feedback participants received after each trial during training and the dimensions on which the stimuli varied. During training, for positive trials (marked + in Table 1) feedback consisted of the phrase “Allergic reaction” in red, followed in the next line by the phrase “10 points of allergy over a total of 35.” On negative trials, feedback consisted of the phrase “No allergic reaction” in black, followed in the next line by the phrase “0 points of allergy over a total of 35.”

During both phases participants were presented at the bottom of the screen with 36 circles with numbers inside ranging from 0 to 35. The left side of the scale was labeled “no allergic reaction” and the right side of the scale was labeled “maximal allergic reaction.” Thus, unlike Experiment 1 in which participants had to predict the allergy using a binary entry (allergy/no allergy), in Experiment 2 they had to predict the exact level of allergy by entering a particular value on the allergy scale.

## Results and Discussion

A total of 3 participants failed the pre-set criteria and were discarded from the study. The final number of participants per group was *n*_*intra*_ = 17, *n*_*extra*_ = 18. Figure 4 shows the results obtained in Experiment 2. Although summation was lower than in Experiment 1, both groups reported higher scores for the compound than the individual cues (F_2, 66_=71.39, p<.01, *η*^2^=.68, 90% CI [.57, .75]), but no detectable difference between groups (F_1, 33_=0.35, p=.56, *η*^2^=.01, 90% CI [.00, .12]) was found. The summation was similar between the groups (F2, 66= 0.71, p=.49, *η*^2^=.02, 90% CI [.00, .09]). Post hoc analysis revealed significant differences for cues A and AB (p<.01) and cues B and AB (p<.01).

**Figure 4.**
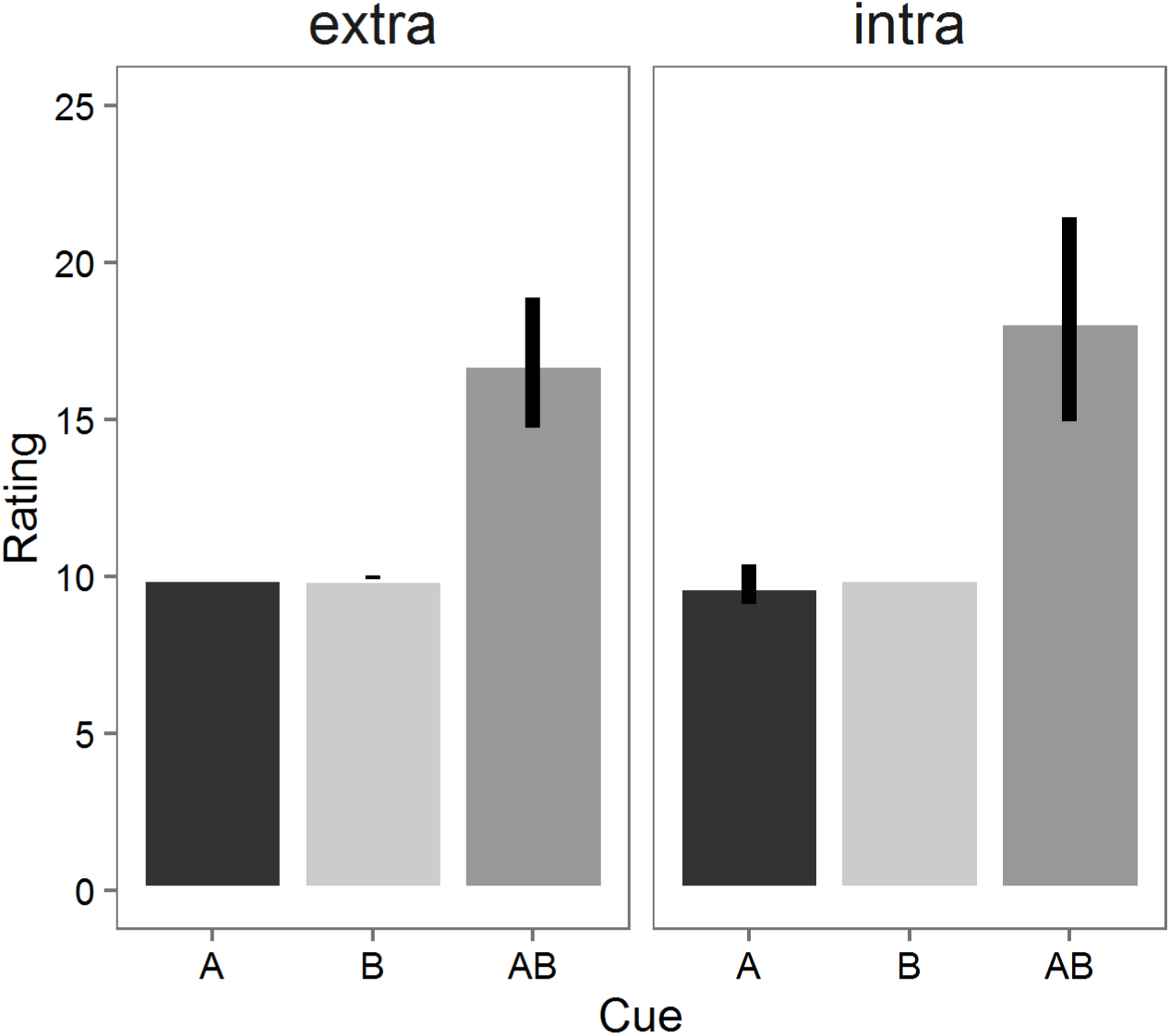
Mean Causal ratings for cues A, B and AB for groups *extra* and *intra* in Experiment 2. Error bars represent 95% confidence intervals.

Experiment 2 shows a similar pattern to that of Experiment 1. Participants in both groups predicted a higher outcome to the compound AB than to each of its constituent elements, but no detectable difference was found between the two groups in terms of the effect. Contrary to the predictions of associative theory, the mean for group intra was higher than the mean for group *extra* in this experiment. This result was obtained regardless of our attempt to increase the difference in similarity between groups *extra* and *intra* by changing the dimensions in which the components A and B varied, and regardless of the change in the rating scale employed to assess the predictions during both phases. The present study is therefore in line with the previous experiment in that summation appears to be a strong effect in humans; the effect, however, does not seem to depend on the rating scale used or the similarity of the cues forming the compound.

## Experiment 3

Taken together, the previous experiments suggest that, if there is an effect of stimulus similarity on summation in human causal learning, then this effect is too small to be detected by our manipulations of brightness, color, shape and rotation. Furthermore, the summation effect appears to be strong even when configural strategies are encouraged by presenting stimuli with a high degree of spatial overlap. Still, the previous experiments did not include a manipulation of spatial distance/overlap as a potential factor determining the level of summation observed. As noted in the introduction, this is a variable that several contemporary models assume to impact on the type of generalization employed (see for example, Soto et al., 2014; Thorwart et al., 2012). In Experiment 1, cues A and B were spatially separated in both groups, and in Experiment 2, cues A and B were spatially contiguous and overlapping in both groups. There was a substantial reduction in the rating to AB from Experiment 1 to Experiment 2 (from an average around 19 to an average of around 17), but it is not clear whether this reduction was due to the change in spatial separation of A and B, or to any of the other changes across experiments (maximum value of the rating scale, stimulus shapes and color, etc.). As the previous literature suggests spatial separation as a potential stimulus factor controlling the level of elemental vs. configural processing of stimuli (see Melchers et al., 2008; Soto et al., 2014a; Thorwart et al., 2012), Experiment 3 was performed to further assess the potential effect of spatial separation of cues on the summation effect.

The same stimuli of Experiment 2 were used in this experiment. However, now A and B differed only in rotation in group *intra*, whereas they differed in rotation and translation (or spatial position; see Figure 5) in group *extra*. This resulted in the components A and B in the compound AB having a high degree of spatial overlap in the group *intra*, but no overlap in group *extra*. A high degree of overlap between the components should have promoted configural encoding in group *intra*. In contrast, spatially-distant stimuli in group *extra* should have promoted elemental encoding. Under these conditions, we expected the summation in group *extra* to be stronger than in group *intra*.

**Figure 5.**
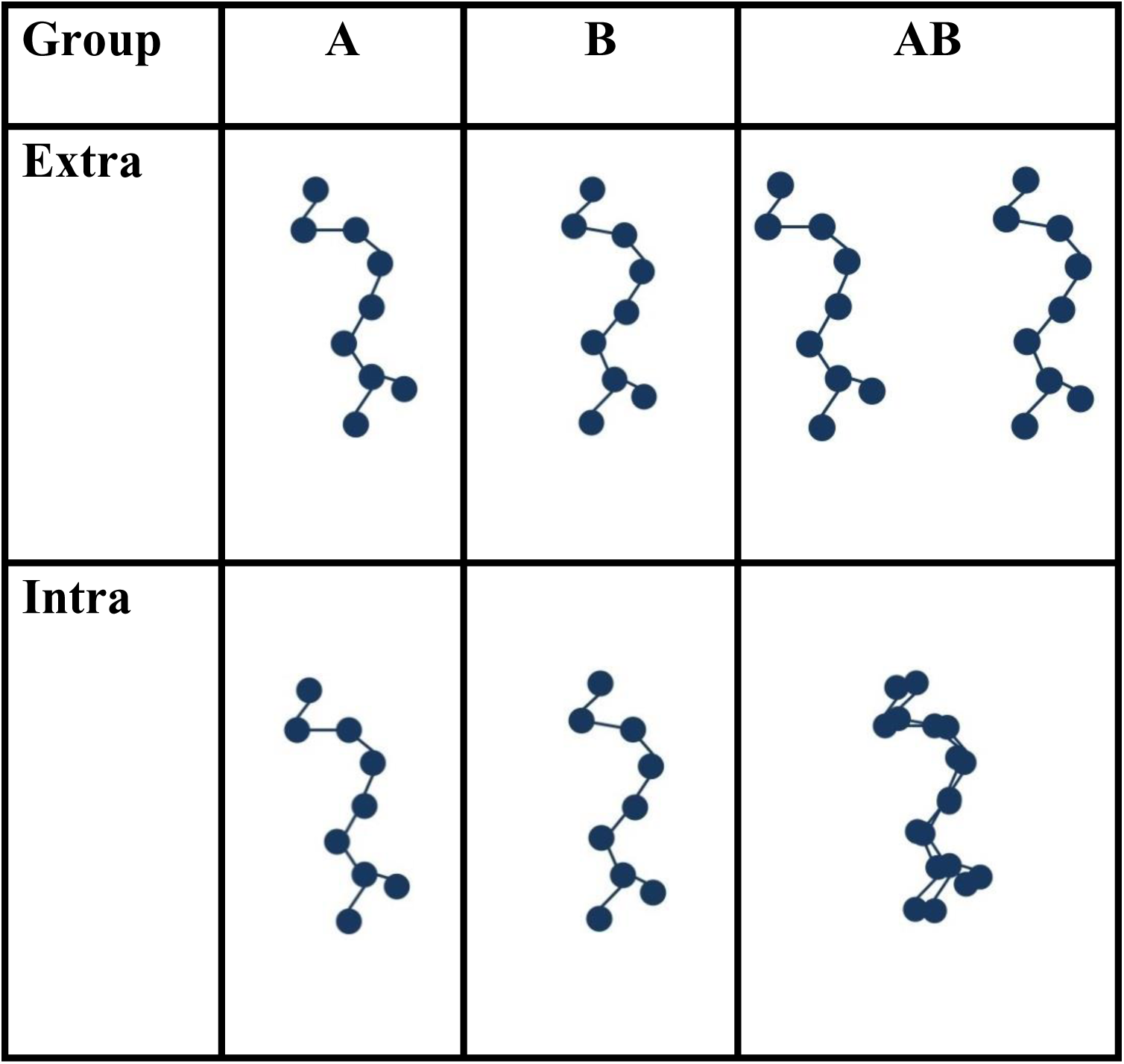
Target stimuli used in Experiment 3. The position of stimuli A and B in the compound AB was counterbalanced in training. For group intradimensional, A and B were presented at the center of the screen; for group extradimensional they were presented to the left and to the right of the center of the screen. A and B respected these positions to form the compound AB during testing.

## Method

### Participants

31 undergraduate students from the University of California, Santa Barbara. Participants were randomly assigned to one of two groups (n_intra_=16, n_extra_=15). They received course credit for their participation in this study.

### Materials

Participants were tested in the same way as in Experiment 2, but cues A and B in group *extra* differed in rotation and spatial location, as shown in Figure 5.

### Procedure

The procedure was the same as in Experiment 2. The stimuli A and B had the same color (dark blue) and shape. For group *intra*, stimulus B had a slight rotation to the right with respect to stimulus A (see Figure 5), but both of them were presented in compound at the center of the screen. For group *extra*, B had a slight rotation to the right, and A and B had a substantial horizontal separation when presented in compound AB.

## Results and Discussion

A total of 5 participants failed the pre-set criteria and were discarded from the study. The final number of participants per group was *n*_*intra*_=14, *n*_*extra*_=12. Figure 6 shows the results of Experiment 3.

**Figure 6.**
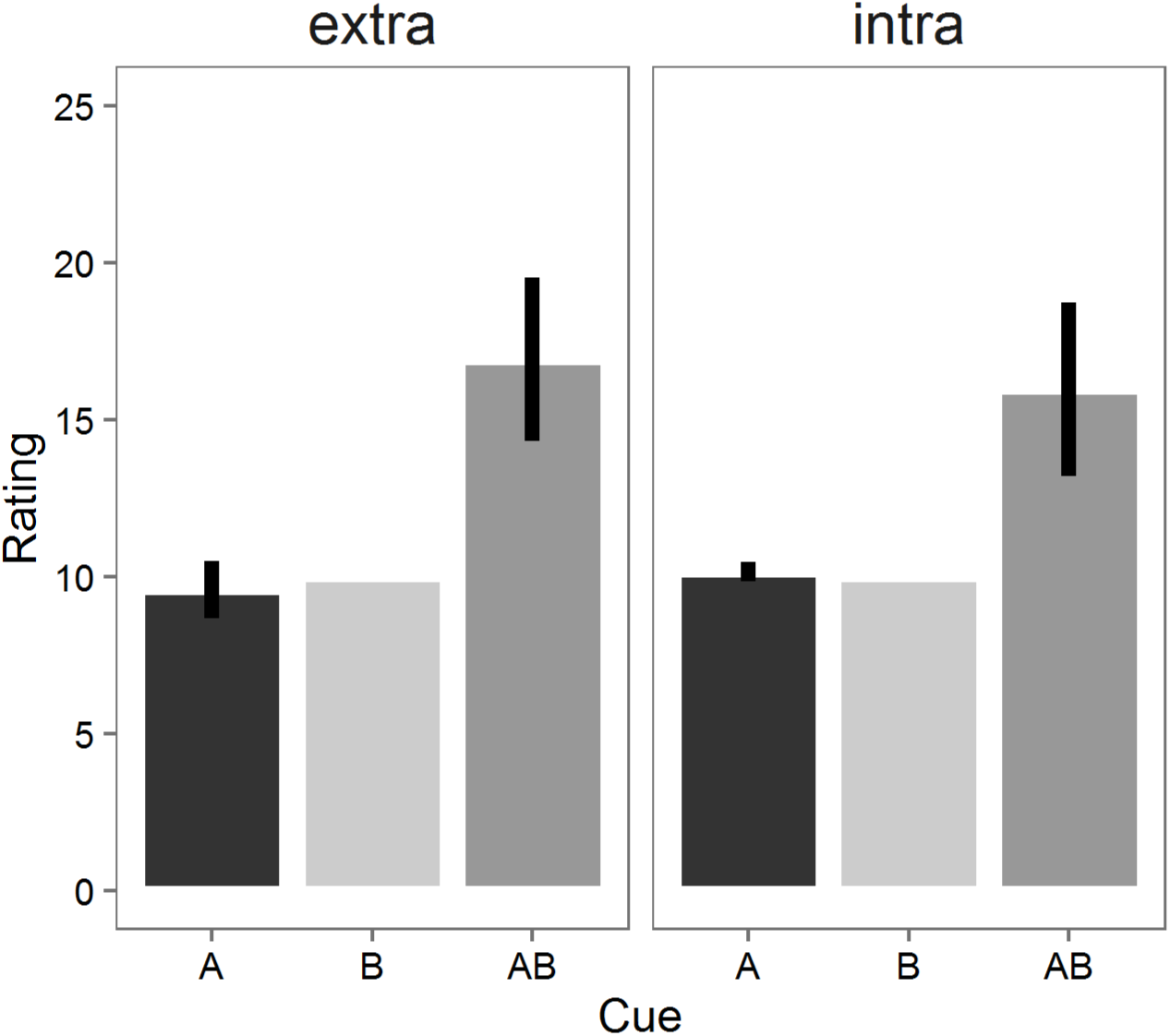
Mean Causal ratings for cues A, B and AB for groups extradimensional and intradimensional in Experiment 3. Error bars represent 95% confidence intervals.

As in previous experiments, groups reported higher scores for the compound than the individual cues (F_2, 48_ =54.41, p<.01, *η*^2^=.69, 90% CI [.55, .76]), but no detectable difference between groups (F_1, 24_ =0.04, p=.41, *η*^2^=.00, 90% CI [.00, .06]) was found. The summation effect did not differ between the groups, as revealed by a nonsignificant interaction between group and cue (F_2, 48_ = 0.56, p=.57, *η*^2^ = .02, 90% CI [.00, .10]).

The pattern of the two previous experiments was replicated in this experiment, with no effect of stimulus properties on the summation effect, regardless of using a different manipulation–spatial contiguity–thought to influence elemental/configural encoding. As spatial contiguity can be construed as an aspect of cue similarity (Harris & Livesey, 2010; Soto, Gershman, & Niv, 2014; Thorwart, Livesey, & Harris, 2012), this experiment aligns with the previous two in showing evidence that the similarity of the cues A and B does not affect the summation effect in human causal learning. The next experiment intended to maximize the effect of stimulus properties by designing A and B in each group so that perceptual similarity was as low as possible in group *extra* and as high as possible in group *intra*.

## Experiment 4

In this experiment, group *intra* was trained with stimuli designed to have only a minimal difference in one dimension, while group *extra* was trained with stimuli designed to have extreme differences in many dimensions. These manipulations should test whether maximizing the stimulus dissimilarity in group *extra*, while minimizing it in group *intra*, can help modulate the summation effect between groups. Using highly dissimilar stimuli should promote elemental encoding and bring about a higher summation effect in group *extra* compared with group *intra*.

## Method

### Participants

72 undergraduate students from the University of California, Santa Barbara. Participants were randomly assigned to one of two groups (n=36 each). They were tested in the same way as in previous experiments and received course credit for their participation.

### Materials

The materials were the same as in previous experiments. For group intra, stimuli A and B varied only slightly in rotation, whereas for group *extra* they had extreme differences in rotation (the object’s main axes were orthogonal to one another), color (dark blue vs. red), spatial position and configuration (see Figure 7).

**Figure 7.**
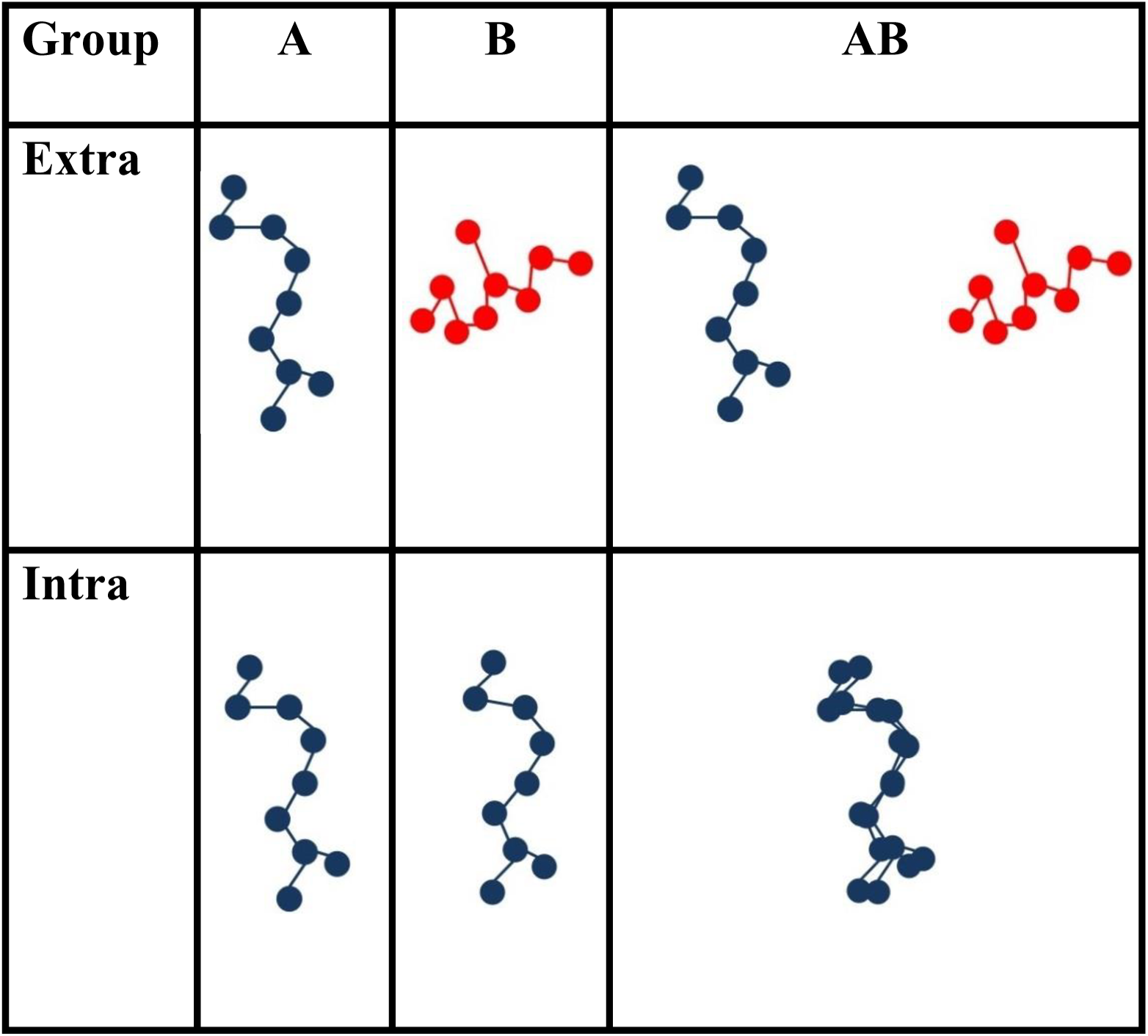
Target stimuli used in Experiment 4. The position of stimuli A and B in the compound AB was counterbalanced in training. For group intradimensional, A and B were presented at the center of the screen; for group extradimensional they were presented to the left and to the right of the center. A and B respected these positions to form the compound AB during testing.

### Procedure

The procedure was the same as in previous experiments.

## Results and discussion

A total of 23 participants failed the pre-set criteria and were discarded from the analysis. The final number of participants per group was *n*_*intra*_=35, *n*_*extra*_=14. Figure 8 presents the results for Experiment 5. As in previous experiments, the mean causal rating to the novel compound AB was greater than each of the cues A and B separately.

**Figure 8.**
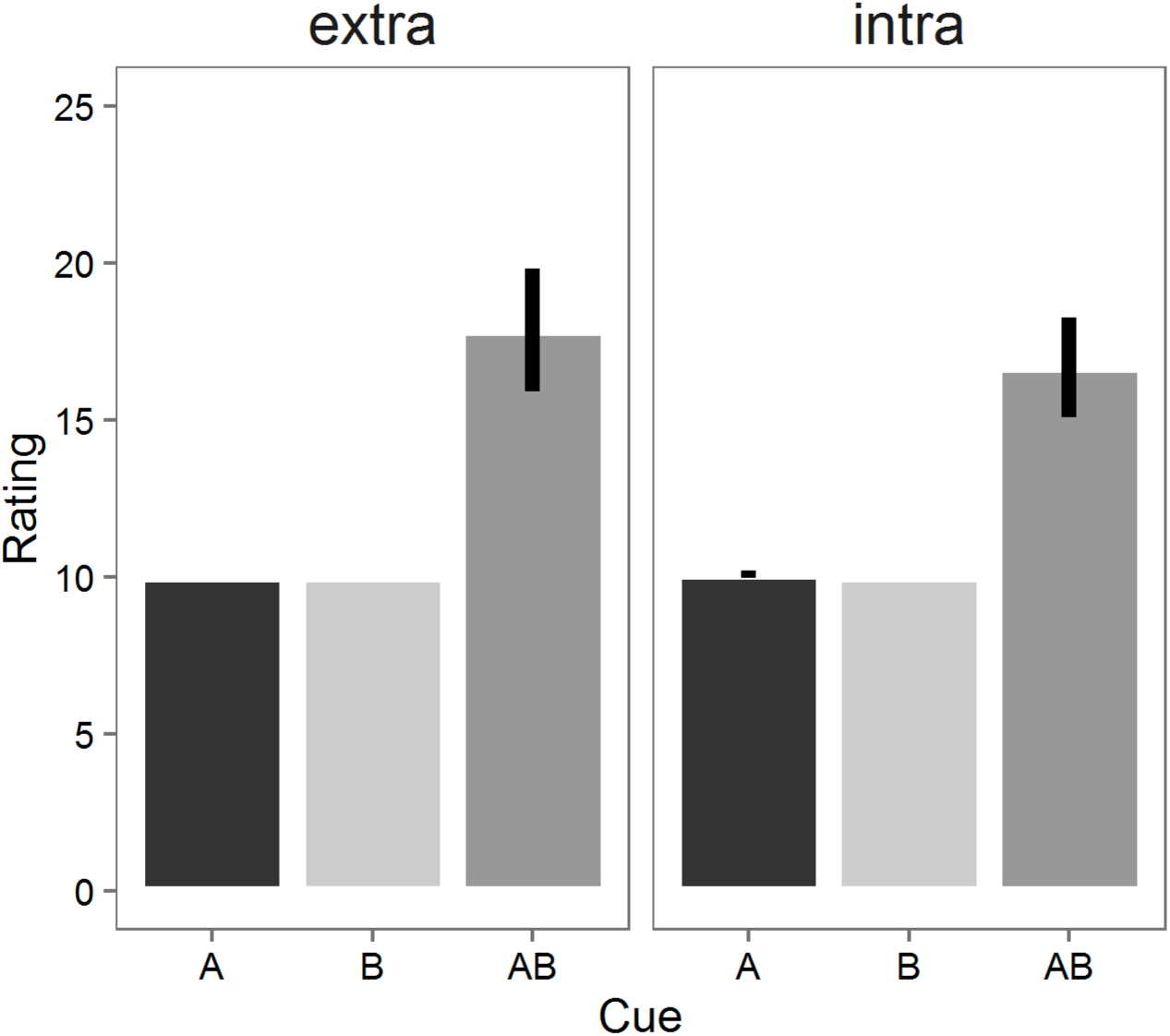
Mean Causal ratings for cues A, B and AB for groups extradimensional and intradimensional in Experiment 4. Error bars represent 95% confidence intervals.

Groups reported higher scores for the compound than the individual cues (F_2, 92_ = 126.47 p<.01, *η*^2^=.73, 90% CI [.65, .78]), but no detectable difference between groups (F_1, 47_ = 0.56, p=.46, *η*^2^=.01, 90% CI [.00, .10]) was found. The summation effect did not differ between the groups (F_2, 92_ = 0.80, p=0.45, *η*^2^=.02, 90% CI [.00, .07]). Post hoc analysis revealed significant differences for cues A and AB (p<.01) and cues B and AB (p<.05).

Results from this experiment reinforce the idea that participants are insensitive to stimulus properties when estimating the causal status of the compound AB. Importantly, the present experiment shows that summation does not differ between groups *intra* and *extra* even when A and B differ in several dimensions and are explicitly designed to maximize elemental processing in group *extra*, while A and B differ only slightly in rotation and are presented with a high degree of overlapping as a compound in group *intra*, a design that should have maximized configural processing.

## Experiment 5

None of the previous experiments found any evidence of an effect of stimulus similarity on the summation effect. However, in all previous experiments the filler cues that predicted no reward, C and D, differed from A and B in many ways (color, shape and configuration). Perhaps stimulus similarity influences the summation effect, but the effect can only be observed when participants are explicitly trained to pay attention to features of A and B that highlight either their similarity or their dissimilarity. Take the stimuli shown in Figure 9. In the row labelled “Intra,” cues A and B differ from cues C and D only in brightness. To successfully categorise these cues into classes that produce and do not produce allergy, people can learn to attend only to stimulus brightness and respond according to a very simple rule: “darker produces allergy”. In the row labelled “Extra”, cues A and B differ from cues C and D in more than just brightness. No single-dimensional rule allows to categorise A and B into one class, and C and D into a different class. There is one darker and one brighter stimulus in each class, and there is one “more circular” and one “more rhomboid” stimulus in each class. To successfully categorise these cues into classes that produce and do not produce allergy, people need to learn the identity of each specific stimulus. This could be done by memorising the value of each stimulus in a single dimension, but is much easier to solve the task by paying attention to differences across both dimensions.

**Figure 9.**
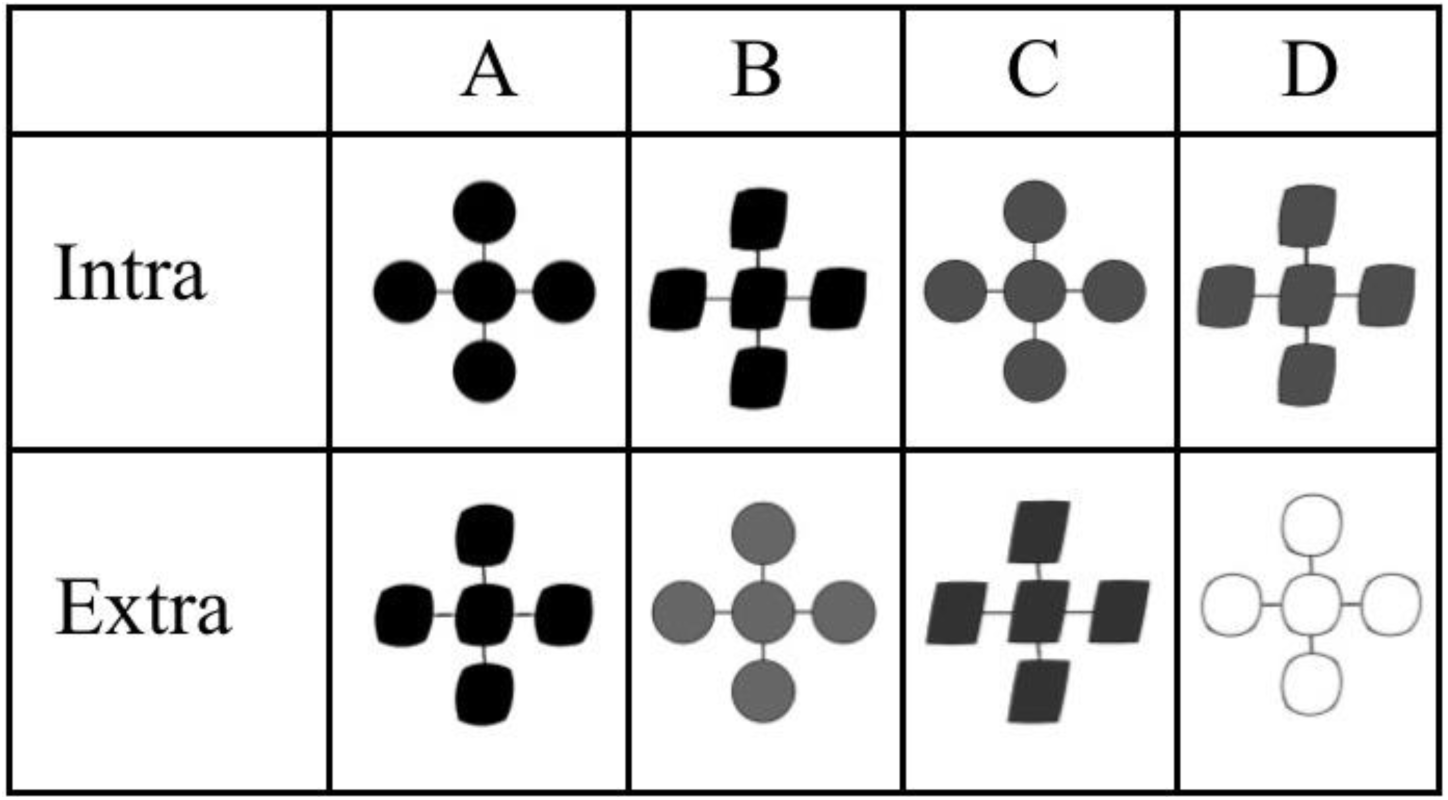
Stimuli used in Experiment 5. In group intra, A and B differed from C and D only in color. In group extra, all stimuli differed both in color and shape.

It should be noted that stimuli in both groups can be described as points in a two-dimensional space created from a shape dimension (circular to rhomboid) and a brightness dimension. The stimuli were selected so as to foster attention to one or two stimulus dimensions, but using similar linearly-separable classification tasks in both groups. While a non-linearly-separable task would have *forced* participants in the *extra* group to pay attention to both dimensions, this would have added a confound to our design. The demands of non-linearly-separable tasks, like negative patterning and biconditional discrimination, are thought to foster configural processing independently of stimulus properties (see Melchers et al., 2008). The goal of this experiment was to promote attention to different stimulus properties, highlighting the similarity or difference between cues A and B and without adding the confound of task demands to our design.

## Method

### Participants

28 undergraduate students from the University of California, Santa Barbara. Participants were randomly assigned to one of two groups (n=14). They were tested in the same way as in previous experiments and they received course credit for their participation.

### Materials

The materials were the same as in previous experiments, but the stimuli used for cues A, B, C and D were as shown in Figure 9.

### Procedure

The procedure was the same as in all previous experiments. The main difference was that the stimuli used in this experiment required participants to differentiate between target cues and fillers in a discrimination task (see Figure 9). The feedback and scales presented to participants were the same as in the previous experiment.

## Results and discussion

A total of 8 participants failed the pre-set criteria and were discarded from the analysis. The final number of participants per group was *n*_*untra*_=8, *n*_*extra*_=12. Figure 10 shows the main results. Both groups estimated a higher allergy magnitude for the compound AB than to each of its constituent elements A and B.

**Figure 10.**
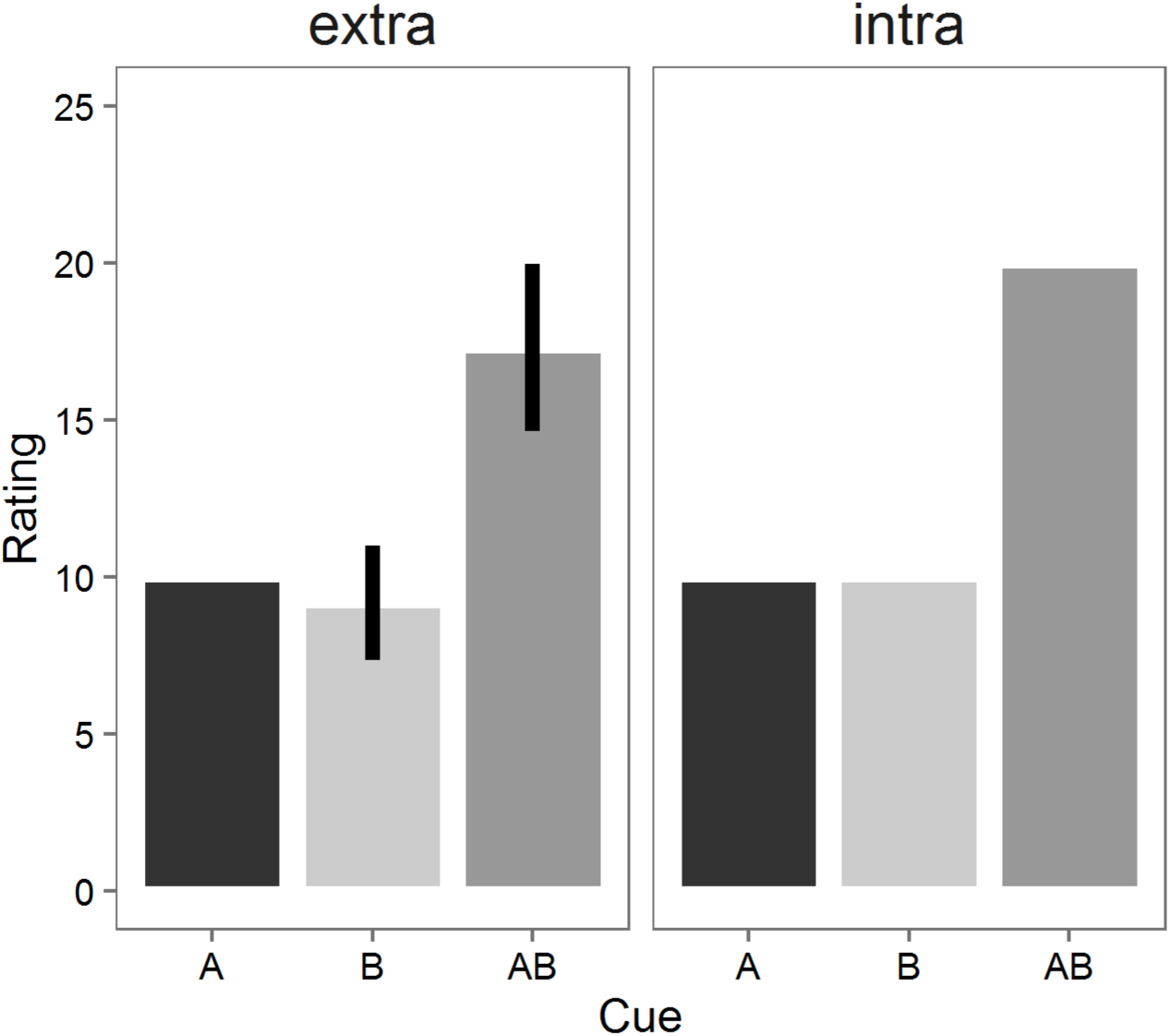
Mean Causal ratings for cues A, B and AB for groups extradimensional and intradimensional in Experiment 5. Error bars represent 95% confidence intervals.

Groups reported higher scores for the compound than the individual cues (F_2, 36_ = 126.7, p= .00, *η*^2^=.88, 90% CI [.80, .91]), but no detectable difference between groups (F_1, 18_ = 2.51, p=.13, *η*^2^=.12, 90% CI [.00, .35]) was found. The summation effect did not differ between the groups (F_2, 36_ = 2.35, p=.11, *η*^2^=.12, 90% CI [.00, .26]). Post hoc analysis revealed significant differences for cues A and AB (p<.01) and cues B and AB (p<.05).

These results show that even when people are trained in a task highlighting either similarities or differences between cues A and B, there is no effect of similarity on the observed summation effect. Even when participants are capable of estimating the levels of the outcome correctly, they nevertheless seem to disregard similarity in forming their predictions.

The results of Experiments 1-5 are largely unfavourable to contemporary associative learning models. As noted in the introduction, most current models designed to explain compound generalization phenomena, such as summation, predict changes in the magnitude of the summation effect when similarity is manipulated, as we have done in all previous experiments. At odds with this prediction, we have found a weak effect of similarity in the difference of the AB scores between the groups *extra* and *intra*. Moreover, the difference obtained was in some experiments numerically opposite to the predictions of similarity-based models.

One could argue that many of these experiments used rather small sample sizes, leading to under-powered statistical tests of our main hypothesis. One way in which these data can be queried through a more powerful test, and provide us with better information as to the size of the effect of similarity in summation is by collapsing all the previous studies in a single estimated statistic. The approach we take here is one suggested by Rouder (2011), who proposes a meta-analytic *t*-test based on Bayesian principles. This method combines the evidence obtained in each experiment in terms of the difference in scores for AB in groups *extra* and *intra* and determines the overall evidence in favor of the null in Experiments 1-5 by calculating a Bayes Factor (*BF*_01_) that compares evidence for the null hypothesis of no difference against the alternative hypothesis of there being a difference in summation between the groups. In our case, the value of *BF*_01_ for Experiments 1-5 was 7.34, implying that the data are 7.34 times more likely to be obtained if there was no difference in summation between the groups than if there was a difference between them. According to Jeffreys (1961), this value can be interpreted as giving substantial evidence for the hypothesis of no difference in summation between groups.

## Experiment 6

One of the central questions in contemporary learning theory is whether the processes underlying many of the most widely-observed phenomena in animal and human learning can be explained by rational principles rather than associative mechanisms (Lovibond & Shanks, 2002; Mitchell, De Houwer, & Lovibond, 2009; Shanks, 2010). For example, rational theory can offer a non-associative explanation for the summation effect by assuming that participants rely on simple prior assumptions regarding the causal independence of A and B in producing the outcome. If A and B are assumed to be independent, the predicted magnitude of an outcome when AB is presented would follow an additive rule similar to that present in elemental associative theory. This independence assumption is, in fact, one of the critical features of several of the most influential rational models of causal induction (Cheng, 1997; Cheng & Novick, 1992; Holyoak & Cheng, 2011b), and a notion that has also been adopted and discussed extensively in contemporary approaches to learning that attribute all associative phenomena as arising from propositional rules (Declercq, De Houwer, &

Baeyens, 2008; Lovibond & Shanks, 2002; Mitchell et al., 2009). For example, the influential Power PC theory proposed by Cheng and colleagues (Cheng, 1997; Holyoak & Cheng, 2011) assumes that the joint predictive power of two candidate causes that independently predict an outcome is given by the noisy-OR rule. If *q*_*A*_ and *q*_*B*_ represent, respectively, the probability of observing the outcome when A or B are present, the noisy-OR rule states that the joint predictive power of the candidate causes A and B will be given by *q*_*A*_ + *q*_*B*_ – *q*_A_*q*_*B*_. Because *q*_*A*_ and *q*_*B*_ are both less than one, the theory predicts a higher causal power -or summation– for the combination of the cues than for each of the cues separately.

From the perspective of rational theory, there is no reason to expect stimulus properties to have an influence on the summation effect. As long as participants rely on the causal independence assumption, they should show the same level of summation across changes in stimulus similarity, as found in Experiments 1-5. The ratings during the test stage would reflect this assumption and be close to a linear sum of each of their single predictions, provided an equal weight to each of the candidate causes is assigned (“A and B are independent, therefore A and B together should cause near 20 points of allergy”).

If the assumption of independence is the main mechanism driving summation in human causal learning, then manipulations other than changes in stimulus properties are likely to bring about rational rules that may impact on the level of summation obtained. Previous research shows, for example, that the application of these type of rules can influence compound generalization. Shanks and Darby (1998) gave participants training with cues A and B followed by the outcome in isolation (A+, B+), but not in compound (AB-), and cues C and D followed by the outcome in compound (CD+), but not in isolation (C-, D-). The critical manipulation was to present participants with the target cues W and X followed by the outcome (W+, X+), and cues Y and Z not followed by the outcome (Y-, Z-). Associative theories would predict that the compound WX will be a stronger predictor of the outcome than YZ, since the latter two have never been followed by the outcome. Surprisingly, this is not what Shanks and Darby (1998) found: participants predicted that the compound WX would be less likely to be followed by the outcome. This is, of course, in sharp contrast with the predictions of any standard associative model, which only considers the associative strength acquired in training by Y, Z, W and X, independently of the relationship between other cues (e.g., A and B) and their compounds (AB). In the Shanks and Darby’s study, however, participants appear to have formed a general rule that made them believe that the relationship between A, B, and C, D with their respective compounds AB and CD during training (“the compound predicts an outcome opposite of that predicted by the individual cues”) should be the same as that of W, X, and Y, Z and their respective compounds WX and YZ during testing.

The aim of this experiment was to explore to what extent weakening summation by manipulations of a simple rational rule of causal independence could bring about differences in summation based on stimulus similarity. To this end, participants were presented during training with examples of cues which predicted the same level of allergy both when presented as a single stimulus and in compound (X+/XX+), and others that predicted no allergy both when presented as a single stimulus and in compound (Y-/YY-). This manipulation prompted participants to learn the rule that that extremely similar (i.e. identical in all aspects, except spatial position) cues combine according to a non-additive rule. In addition, we included the same stimulus manipulation as in Experiment 1 for groups *intra* and *extra*, to determine whether stimulus similarity might have an effect on the summation effect when people assume a non-additive combination rule for extremely similar stimuli. That is, we expected that the non-additive combination rule, learned from extremely similar stimuli, would be applied more readily to similar stimuli in group *intra*, rather than to dissimilar stimuli in group *extra*. Table 3 shows the design of Experiment 6. The only difference with the previous experiments was the inclusion of trials involving the two additional filler cues, X and Y.

**Table 3.**
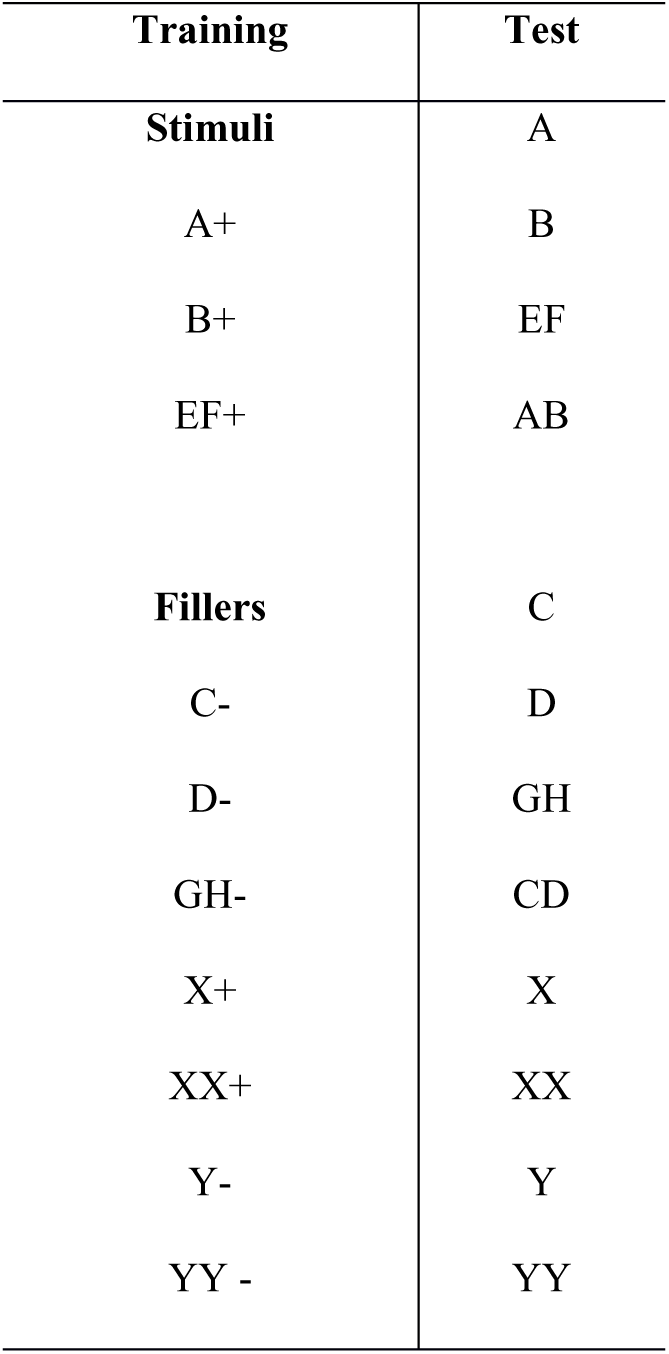
Design of Experiment 6.

## Method

### Participants

46 undergraduate students from the University of Santiago. Participants were randomly assigned to one of two groups (*n*=23). Participants were tested in the same way as in Experiment 1 and they received course credit for their participation.

### Materials

The materials were the same as in previous experiments.

### Procedure

The same stimuli as in Experiment 1 were used. The only difference with Experiment 1 was the addition of the two new fillers, X and Y. Cue X predicted an outcome of 10 points of allergy both when presented as a single stimulus (X+) and in compound (XX+). Cue Y predicted 0 points of allergy both when single (Y-) and in compound (YY-). Participants completed 25 blocks of training, each consisting of a single presentation of each of the 10 trial types shown in Table 3. Trial types were presented in random order within each block.

The total number of training trials was 200. During training and testing we presented participants with the same rating scale as in Experiments 2 and 3 (i.e. a 0-35 points allergy scale).

## Results and discussion

A total of 5 participants failed the pre-set criteria and were discarded from the analysis. The final number of participants per group was *n*_*intra*_=19, *n*_*extra*_=22. Figure 2 shows the results from the main summation test.

Groups reported higher scores for the compound than the individual cues (F_2, 78_ = 9.89, p<.01, *η*^2^=.20, 90% CI [.07, .31]), but no detectable difference between groups (F_1, 39_ = 0.00, p=.93, *η*^2^=.00, 90% CI [.00, .00]) was found. The summation effect did not differ between the groups, as revealed by a nonsignificant interaction between group and cue (F_2, 78_ = 0.021, p=.98, *η*^2^=.00, 90% CI [.00, .01]). Post hoc analysis revealed significant differences for cues A and AB (p<.01) and cues B and AB (p<.05). As in Experiment 1, we also compared the compounds AB and EF. Participants scored AB higher than EF (group intra: mean difference=1.62; group extra: mean difference=1.75). All participants in both groups scored the compound EF as producing 10 points of allergy.

Consistent with our prediction that fostering the adoption of a non-linear combination rule would modulate summation, this effect seemed much weaker for both groups in this experiment. This observation was supported by a comparison of the estimated size of the summation effect for the present experiment and a meta-analytic estimate of the size of the summation effect for all previous experiments. Figure 12 presents a forest plot for Cohen’s *D* (and 95% CI) for the summation effect observed in Experiments 1-5, calculated as the difference between the ratings for the compound AB and the average rating for A and B, collapsing across groups. The overall effect size for the prior five experiments was 1.99, 95% CI [1.57, 2.42], which is four times larger than the effect of 0.5, 95% CI [0.25, 1.14] obtained for the present experiment calculated in the same fashion as in each of the individual studies. The fact that the CI from this experiment did not overlap with the meta-analytic CI from all previous experiments suggests that our manipulation was successful in decreasing the magnitude of summation obtained across both groups.

**Figure 11.**
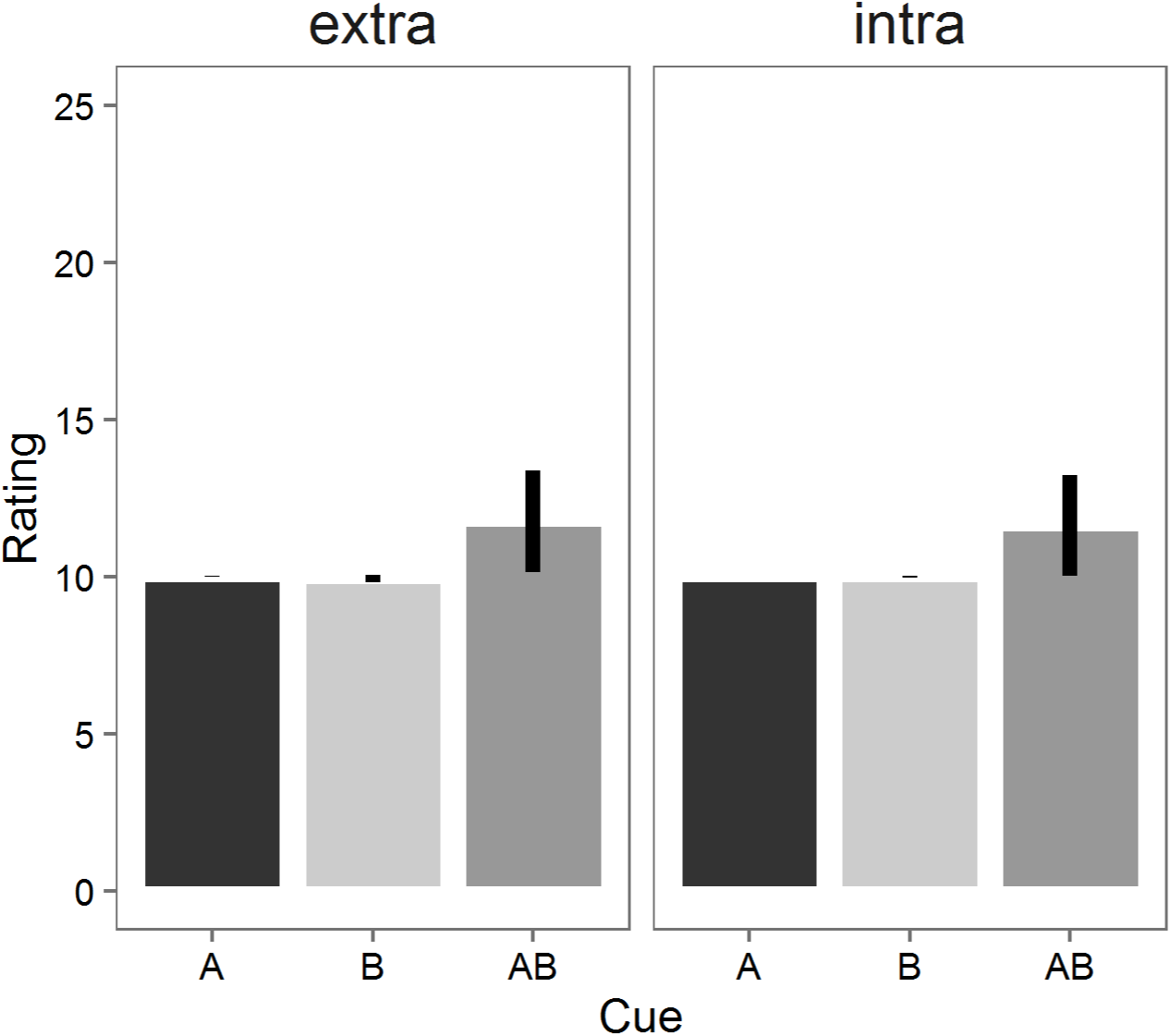
Mean Causal ratings for cues A, B and AB for groups *extra* and *intra* in Experiment 6. Error bars represent 95% confidence intervals.

**Figure 12.**
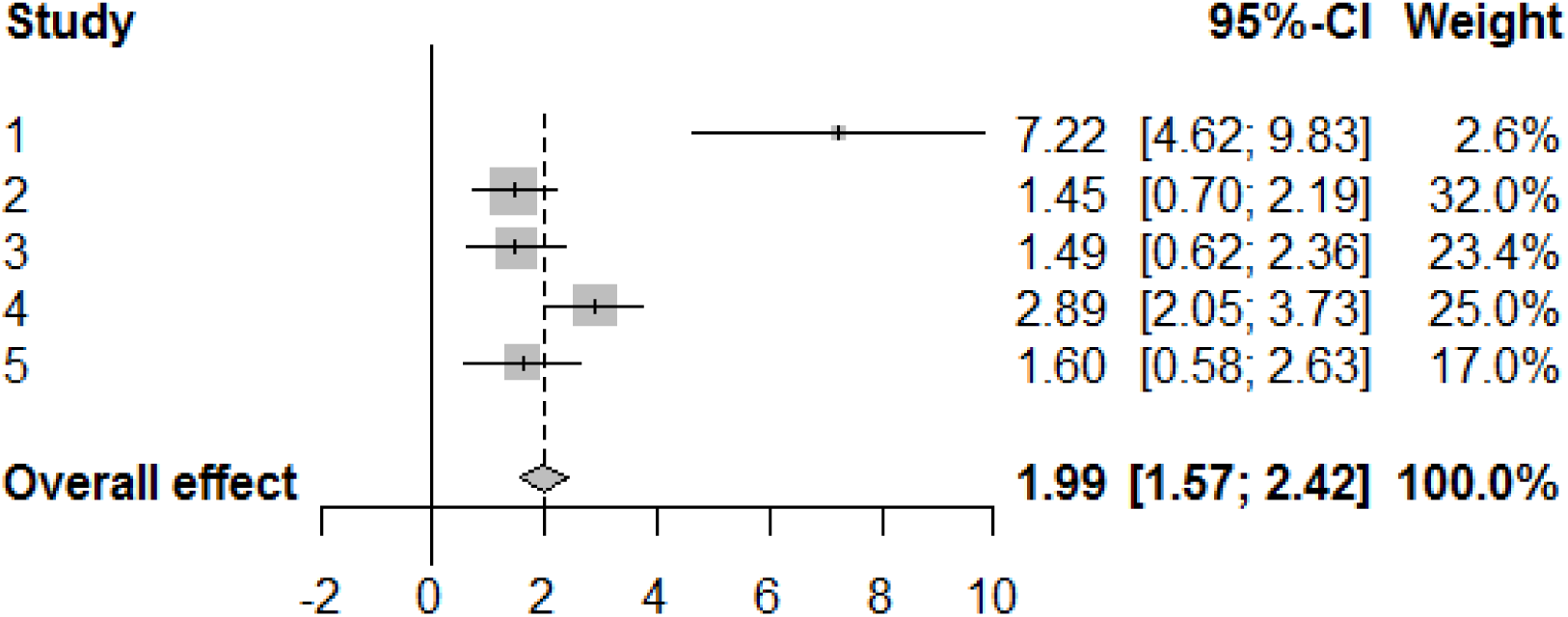
Meta-analysis for the summation effect (score for AB minus score for the average of A and B in both groups) from Experiments 1 to 5. Each dot and error bar represent, respectively, Cohen’s D and 95% CI. The diamond at the bottom represents the overall effect size across all experiments. The size of each dot represents the importance of each study in the overall effect (bigger dots representing higher weighting factors). See Appendix for details.

In sum, a manipulation of stimulus similarity was again ineffective in influencing the summation effect, even when participants were explicitly trained to assume a non-additive combination rule for extremely similar stimuli. Rather than showing a stronger summation effect for more dissimilar stimuli, participants simply applied the same non-additive rule regardless of the properties of cues A and B. These results suggest that a manipulation aimed at modifying participants’ assumption of causal independence successfully influences the magnitude of the summation effect, but does not interact with stimulus similarity.

## General discussion

In the series of experiments reported here, a summation effect was observed across a wide range of experimental manipulations. These results are in accord with previous studies looking into different aspects of summation in causal learning (Collins & Shanks, 2006; Glautier et al., 2010; Soto et al., 2009) and indicate that summation is a robust effect in human causal learning. Manipulating the properties of the stimuli to make them more similar or dissimilar, however, did not produce any difference in the magnitude of summation between the groups.

Experiment 1 demonstrated equivalent summation in groups *intra* and *extra*, in spite of A and B differing only in brightness in the former, while differing both in brightness and shape in the latter. Experiment 2 extended this conclusion, suggesting that the weak effect obtained in Experiment 1 cannot be attributed to features of the rating scale or the spatial distance between components A and B during testing. Furthermore, the fact that changes in rotation and color–rather than shape and brightness–did not change the result, suggests that the lack of an effect of similarity on summation cannot be attributed to the dimensions chosen to manipulate the stimuli. In Experiment 3, spatial separation (Glautier, 2002; Livesey & Boakes, 2004; Melchers et al., 2008; Soto et al., 2014a) was further explored. In group *intra*, A and B were presented at the center of the screen and differed only slightly in rotation but had a high degree of overlapping - a manipulation that should have encouraged configural processing. The components A and B in group *extra* also differed slightly in rotation, but their spatial separation during the test was large, a manipulation that should encourage elemental processing. The results replicated those from the previous two experiments, in that differences in summation failed to emerge even when spatial separation was minimized in group intra, and maximized in group *extra*.

In Experiment 4, the stimuli were designed so that A and B in group *extra* were extremely different, with extreme variations in color and shape. As in Experiment 3, their distance was also maximal on the screen. In contrast, group *intra* was presented with the same stimuli as in Experiment 3. In spite of these manipulations, no detectable difference in summation between the groups was found.

Experiment 5 explored the possibility that the effect of stimulus similarity on summation can only be detected if participants are explicitly trained to pay attention to features of A and B that highlight either their similarity or their dissimilarity. Even though participants were able to correctly predict the outcome after this discrimination task, no differences in summation were found between the groups.

The results from these five experiments are largely unfavourable to associative models which assume that perceptual similarity should affect summation through its impact on the type of generalization strategy adopted by subjects. To recapitulate, these theories anticipate elemental processing and therefore summation with dissimilar stimuli, but configural processing and lower summation with more similar stimuli (Harris & Livesey, 2010; Soto et al., 2014, 2015; Thorwart et al., 2012; Wagner, 2003). None of these predictions were confirmed in the present studies, suggesting that similarity-based models cannot be directly applied to summation in human causal learning.

The fact that we could not detect an effect of similarity on summation prompted us to hypothesise that factors other than stimulus properties needed to be explored. We thought that a reasonable interpretation for these data could be found in rational inductive theories (Cheng, 1997; Cheng & Novick, 1992; Mitchell et al., 2009) and propositional approaches to associative learning (Lovibond et al., 2003; Mitchell et al., 2009; Weidemann, Satkunarajah, & Lovibond, 2016). These theories regard summation as a consequence of a rational account in which participants’ causal attribution to the cues forming a compound is based on a statistical independence assumption. A simple rule that takes each of A and B as a single entity that independently predicts the outcome (for example, the noisy-OR rule proposed by Cheng and colleagues (1997) can account for the fact that summation was consistently obtained in spite of several manipulation of stimulus properties and procedural variables. The main goal of Experiment 6 was thus to weaken this causal independence assumption, thereby testing the extent to which such reduction would allow us to observe differences in summation across the two groups. To this end, stimuli were designed so that identity (the maximum level of similarity possible) did not necessarily yield a higher outcome in compound than each of the elements in isolation. This modification was successful in decreasing summation compared to previous experiments, suggesting that the addition of X+/XX+ and Y-/ YY-trials made participants to form a rule which implied that the relation between A and AB, and B and AB, was similar to that of X and XX or that of Y and YY. The rule, however, was equally applied by both groups, irrespective of the level of similarity of the components A and B. These data indicate that similarity does not affect summation even when rational assumptions of causal independence are minimized.

The main goal of Experiment 6 was to test whether weakening the causal independence assumption would produce an effect of similarity on summation, rather than testing whether rational rules were driving summation. Still, when the results from this experiment are compared to those from all the other experiments in our study, they suggest that, as in the case of learning phenomena such as blocking (Beckers, De Houwer, Pineno, & Miller, 2005; Lovibond et al., 2003), the phenomenon of generalization – which takes place after the learning stage - can also be affected by manipulations of assumptions about independence. The evidence for this claim is only suggestive, as it is based on a between-experiments comparison. Ideally, one would like to compare summation in two groups that only differ with respect to the outcomes produced by XX and YY.

The observation of summation being affected by pretraining with unrelated cues can be explained by recent models based on Bayesian principles (Holyoak & Cheng, 2011; Lu, Rojas, Beckers, & Yuille, 2016; Lucas & Griffiths, 2010). These models are predicated under the assumption that in a causal learning task subjects weigh different hypothesis about the underlying causal structure of the world. Using Bayes’ rule during training, subjects are able to update possible hypothesis about the form of the relationship between the compound AB and the outcome. This flexibility allows these models to explain why a disjunctive hypothesis such as that found in the noisy-OR rule may be modified to reflect non-linearity as subjects observe instances of other cues which violate this prior assumption of independence (Lu et al., 2016).

If indeed participants in all previous experiments were adopting a rational rule based on the noisy-OR rule alone, the rating for AB in Experiment 6 should have been similar to that of A or B in isolation. This was not, however, what we found: although much weaker than in all previous experiments, the summation effect was still present in both groups. This raises the possibility that an associative process was still playing a role. A rational analysis would suggest that the fact that summation is still observed is a consequence of some participants simply not adjusting to the non-additivity examples provided in training. If this is so, then those participants that did not adopt the rational rule should have scored the compound as 20; the rest should have adopted the rule and scored the compound as 10. One way of exploring this possibility is to generate a more detailed visualization of the individual data for this experiment. Figure 13 shows the individual summation scores (AB – mean of A and B) in both groups in Experiment 6. As can be appreciated, the majority of participants in each group has a summation score of 0, implying that they scored the compound as producing the same outcome as the mean of the individual cues. In addition, only 2 participants in each group show a score of 10, indicating that they did not adopt the rational rule and instead showed a similar pattern to the previous experiments. Seven participants showed summation scores in between 0 and 10. These data show that the majority of people adopted a rational rule. The fact that there is a summation effect in this experiment is therefore explained by the fact that some participants scored the compound AB higher than A and B. However, there is no evidence of a group of participants consistently using a linear additive rule as in the previous experiments reported here. What is clear from this analysis is that there was a significant number of participants scoring AB as producing the same level of allergy as A and B on average.

**Figure 13.**
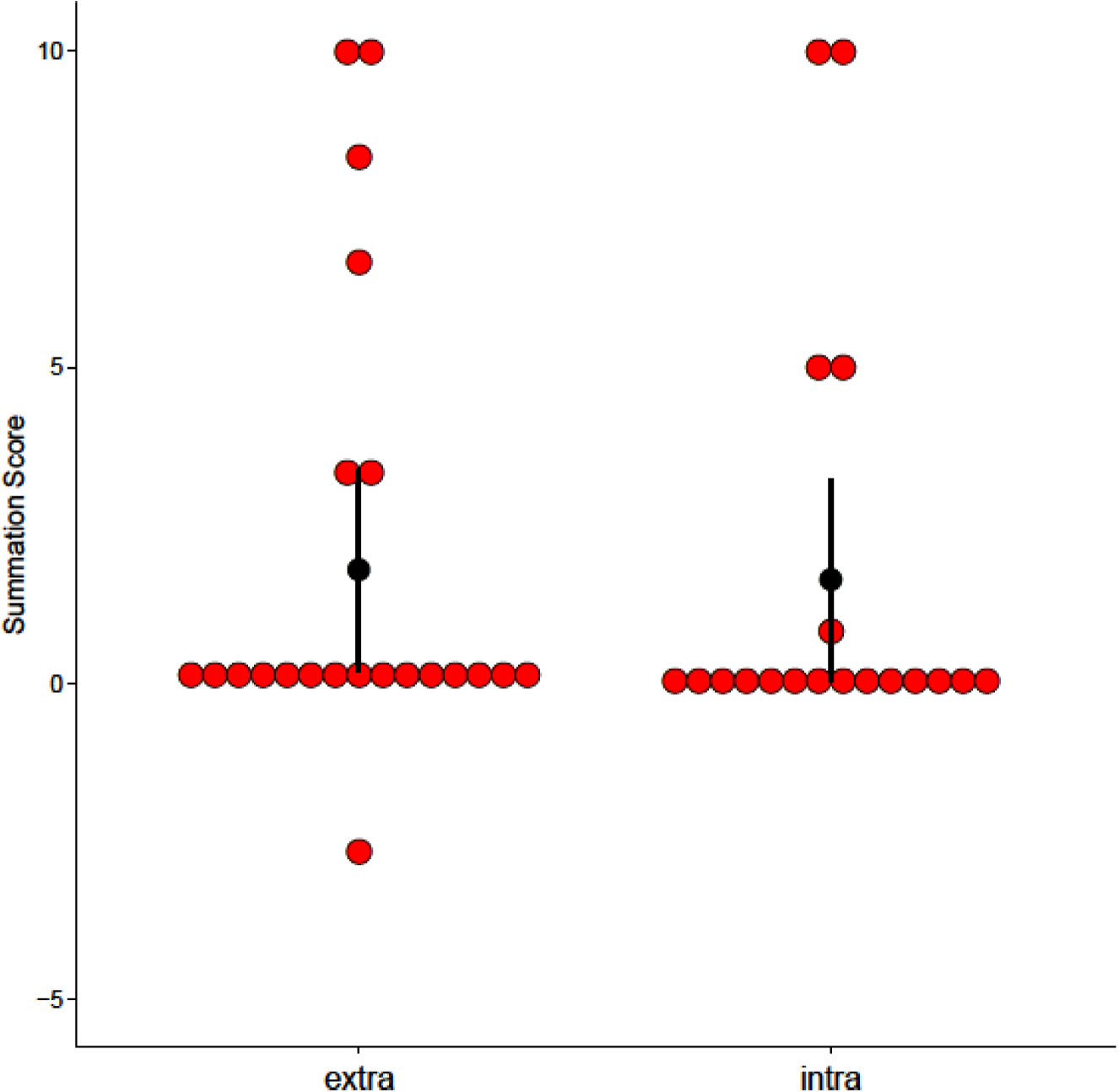
Summation scores (Score for AB - mean score of A and B) for groups extra and intra in Experiment 6. The red dots indicate individual data points per subject. The black dot and error bar indicate, respectively, mean and 95% CI.

In spite of not finding evidence of similarity affecting summation, the importance of stimulus features cannot be completely ruled out from these experiments. It may well be the case that the role of stimulus features is to provide support for rational assumptions. An associative perspective, for example, would interpret spatial separation as a factor that promotes elemental processing and therefore higher summation during test. Livesey and Boakes (2004) found this to be the case for blocking: cues far apart on the screen produced blocking, whereas spatially-contiguous cues did not. However, it could equally be argued that spatially-separated cues are prompting participants to assume these are causally independent, whereas spatially-contiguous cues could lead them to believe they are not.

One possibility by which similarity-based models could accommodate these data is assuming that participants in the intra groups were not able to discriminate at all between A and B. This implies that a compound AB should be regarded as AA+ (or BB+) during training. It could be argued that in such a case no perceptual interaction between cues occurs, and therefore summation is predicted for AA during the test. Contemporary models such as Wagner’s REM (2003) do not discuss this possibility, but an argument could be made that in such a case no replacement should occur. However, it is difficult to reconcile the idea that an AA trial, in which two identical stimuli are presented, should be similar to one including two very dissimilar stimuli in terms of replacement of elements. Other similarity-based models, like Soto et al. (2014)’s, do include spatial separation as an aspect of similarity precisely to deal with these extreme cases in a more explicit manner: an AA trial should be considered as one including two extremely similar stimuli that only differ in spatial position.

Although the application of a noisy-OR rule seems sufficient to explain these data, it is possible that participants would deploy the rule only to the extent that stimulus features do not undermine the independence assumption. The obvious possibility is that the rule should be applied to the extent that participants are able to distinguish A and B in the compound AB - if there is no perceptual interaction between them, then there is no need for similarity-based processes to be deployed. Perhaps employing more complex compounds or generalization tasks - such as, for example, those in which the generalization goes from compounds to elements - would make participants rely on similarity at the expense of rational rules.

The main conclusion that can be drawn from this series of experiments is the absence of an effect of similarity on summation when employing unimodal visual stimuli. Participants in a simple allergy task (Lovibond et al., 2003; Soto et al., 2009; Van Hamme & Wasserman, 1994) seem to rely on simple linear additive rules based on causal independence, and tend to disregard similarity in assessing their predictions. Our results are thus at variance with those reported by Redhead (2007), who, as in the Pavlovian conditioning case, found that the summation effect was affected by whether the compound was comprised of unimodal or multimodal components - when using multimodal components, summation was obtained; when using unimodal, it was not. In contrast, we found that varying degrees of similarity in unimodal visual stimuli does not play a role in modulating summation. Although it remains to be tested whether these results can be extended to other type of scenarios such as fear conditioning and predictive learning, it is clear that, at least under a simple allergy task and a wide range of conditions, similarity is not a factor in modulating the summation effect in causal learning.

## Acknowledgements

We thank Julián Contreras for helping with data collection for some of the experiments reported in this paper.

## Appendix

The two meta-analyses reported here were carried out using R version 3.3.0 and RStudio version 1.0.136 (RStudio Team, 2015), extended with the *BayesFactor* 0.9.12 (Morey & Rouder, 2015) and *meta* 4.8.3 (Schwarzer, 2007) packages.

### Meta-analysis 1: Bayesian meta-analytic t-test.

For the first analysis, we obtained the difference between the mean scores for the AB compound between groups *extra* and *intra* and calculated an independent t-test for Experiments 1-5. These values were then used, together with the sample sizes for groups *intra* and *extra* on each study, to obtain an overall Bayesian Factor (*BF*_01_) for these studies.

### Meta-analysis 2: Overall summation effect.

Our selected effect size was Cohen’s *D*, as it allowed us to standardize the difference between the mean scores obtained in each study. Although it could be argued that calculating a raw difference between means would have sufficed to analyse the overall effect of these experiments, the fact that there were changes in the rating scale presented to participants led us to favor the selection of *D* as our effect size of interest.

Given the similarity of Experiments 1-5 in terms of the methods and question involved, it is reasonable to assume that they all estimate the same underlying effect. In other words, the estimated population effect size can be regarded as fixed, and all the variability observed by randomness between experiments. In meta-analysis terminology, this is known as a *fixed effect model* (Borenstein, Hedges, Higgins, & Rothstein, 2009). This was the method chosen for our analysis.

For the analysis, the overall summation effect, SEF, was calculated as *SEF* = *AB* – *mean*(*A*, *B*), where AB represents the mean score for *AB*, and *mean*(*A*, *B*) the mean of A and B together, collapsed across both groups. The value of D for each of the studies was then calculated as 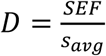, where *s*_*avg*_ represents the average of the standard deviations obtained for *AB* and *mean*(*A.B*), respectively, given by 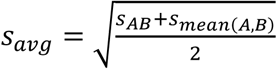 The values obtained where then entered into the *meta* package to obtain the results.

## Supplementary material

Supplementary tables. Mean ratings (and standard errors) for all the fillers tested in each experiment.

**Experiment 1.**

|  | Group | C | D | CD | EF | GH |
| --- | --- | --- | --- | --- | --- | --- |
| Rating | extra | 0.00 (0.00) | 0.00 (0.00) | 0.00 (0.00) | 10.00 (0.00) | 0.00 (0.00) |
|  | intra | 0.00 (0.00) | 0.00 (0.00) | 0.00 (0.00) | 10.20 (0.17) | 0.00 (0.00) |

**Experiment 2.**

|  | Group | C | D |
| --- | --- | --- | --- |
| Rating | extra | 0.00 (0.00) | 0.00 (0.00) |
|  | intra | 0.00 (0.00) | 0.00 (0.00) |

**Experiment 3.**
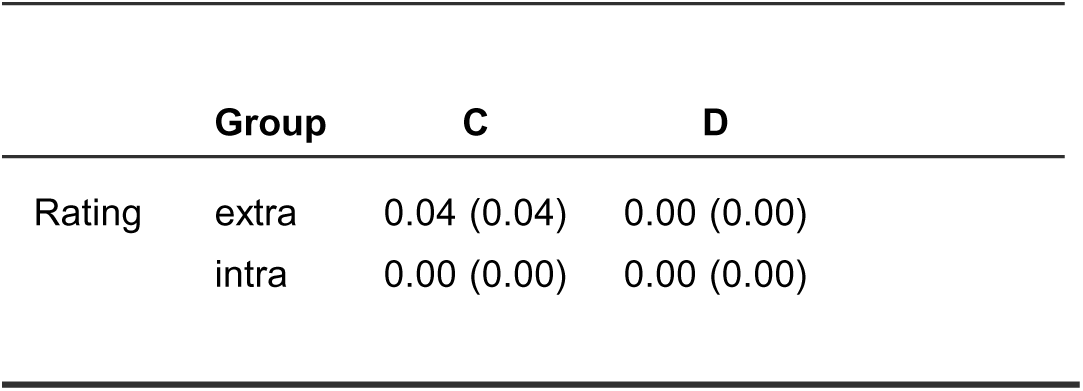

SUMMATION IN CAUSAL LEARNING **Experiment 3**
|  | **Group** | **C** | **D** |
| --- | --- | --- | --- |
| Rating | extra | 0.04 (0.04) | 0.00 (0.00) |
|  | intra | 0.00 (0.00) | 0.00 (0.00) |

**Experiment 4.**

|  | **Group** | **C** | **D** |
| --- | --- | --- | --- |
| Rating | extra | 0.00 (0.00) | 0.00 (0.00) |
|  | intra | 0.00 (0.00) | 0.00 (0.00) |

**Experiment 5.**

SUMMATION IN CAUSAL LEARNING **Experiment 5**
|  | **Group** | **C** | **D** |
| --- | --- | --- | --- |
| Rating | extra | 0.00 (0.00) | 0.00 (0.00) |
|  | intra | 0.83 (0.53) | 0.00 (0.00) |

**Experiment 6.**

|  | **Group** | **C** | **D** | **CD** | **EF** | **GH** | **X** | **XX** | **Y** | **YY** |
| --- | --- | --- | --- | --- | --- | --- | --- | --- | --- | --- |
| Rating | extra | 0.00 (0.00) | 0.00 (0.00) | 0.15 (0.15) | 10.0 (0.00) | 0.08 (0.07) | 10.00 (0.01) | 10.30 (0.30) | 0.00 (0.00) | 0.20 (0.20) |
|  | intra | 0.00 (0.00) | 0.00 (0.00) | 0.00 (0.00) | 10.0 (0.00) | 0.08 (0.08) | 10.00 (0.00) | 10.00 (0.00) | 0.02 (0.01) | 0.00 (0.00) |

## Footnotes

1 Here we include only models that have explicitly performed simulations showing an impact of similarity on summation. Other models, like that of McLaren and Mackintosh (2002), might yield similar predictions under some conditions. However, since they have not so far reported this type of simulations, we did not include them as supporting this assumption.

2 We should note that the terms extra and intra refer to dimensions, not modalities; our task deploys unimodal visual stimuli with different degrees of similarity as measured by configuration, rotation, spatial position, and color. In all the experiments reported here, intra should simply be taken as indicating “more configural processing” while extra indicating “more elemental processing”.

